# Genetic Susceptibility to Multiple Sclerosis: Interactions between Conserved Extended Haplotypes of the MHC and other Susceptibility Regions

**DOI:** 10.1101/603878

**Authors:** DS Goodin, P Khankhanian, PA Gourraud, N Vince

## Abstract

**OBJECTIVE:** To study the accumulation of MS-risk resulting from different combinations of MS-associated conserved-extended-haplotypes of the MHC and three non-MHC risk-loci nearby genes EOMES, ZFP36L1, CLEC16A.

**BACKGROUND:** Defining “genetic-susceptibility” as having a non-zero probability of developing MS, both theoretical considerations and epidemiological observations indicate that only 2.2–4.5% of northern-populations can possibly be “genetically-susceptible” to MS. Nevertheless, many haplotypes (both within the MHC and elsewhere) are unequivocally MS-associated and, yet, have population-frequencies of >20%. Such frequency-disparities underscore the complex-interactions that must occur between these “risk-haplotypes” and MS-susceptibility.

**DESIGN/MEHTODS:** The WTCCC dataset was statistically-phased at the *MHC* and at three other susceptibility-regions. Haplotypes were stratified by their impact on “MS-risk”. MS-associations for different combinations of “risk-haplotypes” were assessed. The appropriateness of both additive and multiplicative risk-accumulation models was determined.

**RESULTS:** Combinations of different “risk-haplotypes” produced an MS-risk that was considerably closer to an additive model than a multiplicative model. Nevertheless, neither of these simple probability-models adequately accounted for the accumulation of disease-risk in MS at these four loci.

**CONCLUSIONS:** “Genetic-susceptibility” to MS seems to depend upon the exact state at each “risk-locus” and upon specific gene-gene combinations across loci. Moreover, “genetic-susceptibility” is both rare in the population and, yet, is a necessary condition for MS to develop in any individual. In this sense, MS is a “genetic” disease. Nevertheless although, “genetic-susceptibility” is a necessary condition for MS to develop, environmental factors (whatever these may be) and stochastic processes are also necessary determinants of whether a “genetically-susceptible” individual will actually get MS.

**Author Summary:** Defining a “genetically-susceptible” individual to be any person in the population who has any chance of developing multiple sclerosis (MS), we demonstrate that, at a theoretical level and using widely-accepted epidemiological observations, only 2.2-4.5% of individuals in northern populations can possibly be “genetically susceptible” to MS. Thus, more than 95.5% of individuals in these populations have no chance of getting MS, regardless of the environmental circumstances that they may experience.

Nevertheless, certain “susceptibility-haplotypes” (e.g., *HLA-DRB1*15:01~DQB1*06:02*) have a far greater carrier-frequency than 2.2-4.5%. Consequently, most carriers of these “susceptibility-haplotypes” have no chance of getting MS and, therefore, their “susceptibility” must arise from some combination of these haplotypes with other “susceptibility-haplotypes”. By analyzing such combinatorial impacts at four susceptibility-loci, we found significant interactions both within and between the different “susceptibility-haplotypes”, thereby confirming the relationship between “genetic-susceptibility” and specific gene-gene combinations.

The nature of “genetic-susceptibility” developed here is applicable to other complex genetic disorders. Indeed, any disease for which the MZ-twin concordance rate is substantially greater than the life-time risk in the general population, only a small fraction of the population can possibly be in the “genetically-susceptible” subset (i.e., have any chance of developing the disease).

## Introduction

The nature of susceptibility to multiple sclerosis (MS) is quite complex and involves both environmental and genetic factors [1-4]. Recently, considerable progress has been made in our understanding of the basis for “genetic-susceptibility” in MS. Thus, to date, over 200 common risk variants (located in diverse autosomal genomic regions) have been identified as being MS-associated by genome-wide association screens (GWAS) using large arrays of single nucleotide polymorphisms (SNPs) scattered throughout the genome [5-14]. Despite this recent explosion in the number of identified MS-associated regions, however, the association of MS susceptibility with certain alleles of the human leukocyte antigens (*HLA*) inside the major histocompatibility complex (*MHC*) has been known for decades [11,15-22]. Also, the importance of these new observations to our understanding of “genetic-susceptibility” in MS is tempered by the fact that any single SNP is generally associated with more than one gene or with more than one allele of a single gene. Moreover, sometimes the presumptively associated (i.e., “candidate”) genes are at a considerable genetic distance from the location of the SNP itself [13,14].

For example, we have recently identified an 11-SNP haplotype (*a1*), which spans 0.25 megabases (mb) of DNA surrounding the *HLA-DRB1* gene on the short arm of chromosome 6, and which has the most significant association with MS of any SNP haplotype in the genome [23,24]. Moreover, 99% of these (*a1*) SNP haplotypes carry the *HLA-DRB1*15:01~HLA-DQB1*06:02* haplotype and, conversely, 99% of these *HLA-*haplotypes carry the (*a1*) SNP haplotype. In the Welcome Trust Case Control Consortium (WTCCC) dataset, the odds ratio (*OR*) for an association the full *HLA-DRB1*15:01~HLA-DQB1*06:02~a1* haplotype was 3.28 (p<<10^-300^) and similar disease associations for portions of this haplotype have been consistently reported in many other studies from northern MS populations [11,15-22,25]. Nevertheless, despite this extremely close association, and despite the fact that many of these 11 SNPs, individually, are highly associated both with this particular *HLA-*haplotype and with MS, for none of these individual SNPs is this association exclusive [26]. Thus, each of these SNPs is also found in association with other *HLA-*haplotypes [24,26]. Consequently, even with the large number of SNPs now identified as being MS-associated [13,14], any such association can only be viewed as simply tagging a relatively large genomic region; it cannot be used with confidence to identify any specific gene or to implicate any specific allele with respect to its role in causing, or contributing to, a “genetic-susceptibility” for MS.

Using data from the WTCCC, we recently reported that the *MHC* region was largely composed of a relatively small collection of highly conserved extended haplotypes (CEHs), stretching across all of the “classical” *HLA* genes (*HLA-A, HLA-C, HLA-B, HLA-DRB1*, and *HLA-DQB1*) – a distance spanning more than 2.7 mb of DNA [26]. As shown in *File S2 (Supplemental Fig D*), this same basic population structure is also found in numerous other widely separated human populations around the world [25]. These CEHs seem to be under a strong selection pressure, presumably based upon favorable biological properties of the complete haplotype [26]. Lastly, this population structure is unlikely to be the result of a linkage disequilibrium caused by the founder effects of a small population migrating out of Africa and radiating throughout Eurasia and the Americas. Rather, the marked divergence of the CEH composition both among and between these different human groups, including Africans (*File S2; Tables S4a & S4b*), indicates that this population structure must be due to selection. Consequently, “genetic-susceptibility” to MS, at least in so far as it relates to the *MHC*, is not likely to be attributable to any specific *HLA* allele but, rather, seems to depend upon the nature of each CEH [26]. Nevertheless, because many CEHs seem to be selected simultaneously and because the exact composition of the selected CEHs seems to be so fluid between different populations, the actual fitness landscape for this selection must be extremely variable in space and/or time and the introduction of novel allelic combinations must occur quite frequently [26].

Indeed, in the mostly European WTCCC population, the most frequent (and, thus, the most highly selected) Class II haplotype is *HLA-DRB1*15:01~HLA-DQB1*06:02~a1*, which accounted for 12.4% of all Class II haplotypes present in the control population. Nevertheless, most (or all) of the CEHs, which contain this Class II *HLA*-haplotype (including those whose full CEH had only a single representation in the WTCCC), are associated with an increased MS-risk, although the magnitude of the association varies significantly among the different CEHs [25]. Moreover, some rare haplotypes, which include the Class II motif of *HLA-DRB1*15:01~HLA-DQB1*06:02* but not (*a1*), seem not to carry any risk [26]. By contrast, haplotypes containing (*a1*), but not this Class II *HLA-*motif, still carry substantial risk [26]. For the Class II *HLA-*motif of *HLA-DRB1*03:01~HLA-DQB1*02:01*, this dependence on the nature of the full CEH was even more evident. Thus, carriers of *HLA-DRB1*03:01~HLA-DQB1*02:01~a2* seem to have a disease risk that is either dominant or dose dependent whereas most carriers of *HLA-DRB1*03:01~HLA-DQB1*02:01~a6*, seem to have a disease risk that is recessive or “neutral” [26]. Nevertheless, at least one such haplotypes (i.e., *HLA-A*\**24:02~C*07:01~HLA-B*08:01~HLA-DRB1*03:01~HLA-DQB1*02:01~a6*) has either a dominant or a dose dependent disease risk [26]. These examples underscore the complex interactions that take place between the various *MHC* alleles/haplotypes and MS-risk.

In the present manuscript, we explore these relationships and interactions between the different disease-associated CEHs in the MHC region and other “risk” haplotypes elsewhere in the genome, in order to shed light on the nature of “genetic-susceptibility” to MS. Before embarking, however, we develop two underlying theoretical considerations. First, we summarize what is currently known about the various epidemiological parameters, which are associated with “genetic-susceptibility” in MS, and, in particular, about those parameters potentially related to the role of the MHC and other loci in producing this susceptibility. Second, we consider the different risk models used in epidemiology to account for the accumulation of disease-risk, which is caused by the combination of one or more MS-risk factors in the same individual.

### Epidemiological Parameters Associated with MS

Several epidemiological parameters, which are associated with MS pathogenesis, can be defined and estimated and the relationships between them established using published epidemiological data. The definitions for the model parameters are provided in Table 1. Thus, population parameters, which are directly observable in the population as a whole, can be used to estimate non-population parameters, which cannot be measured directly, but which are, nonetheless, of considerable theoretical interest (*File S1*). These estimates and these relationships, summarized here, are developed comprehensively in the *S1 File*. For example *P*(*MS*), which represents the life-time probability that an individual in the general population (*Z*) will develop MS can be estimated by three methods (*File S1*) based upon both the distribution of onset-ages for MS and the increased mortality experienced by MS patients [27-33], and each of these methods provides a remarkably consistent estimate for *P*(*MS*) in northern populations, which is ~0.3%.

**Table 1.** Definitions for Epidemiological Parameters used in the Model*

| Parameter | Definition |
| --- | --- |
| $(Z)$ | Set of all individuals in the population |
| $P(MS)$ | Life-time probability of getting *MS* for members of $(Z)$ |
| $(G)$ | Set of all individuals in $(Z)$ who have some chance of getting *MS* |
| $(G-)$ | Set of all individuals in $(Z)$ who have no chance of getting *MS* |
| $(H+)$ | Set of all carriers of the *DRB1\*15:01~DQB1\*06:02~a1* haplotype in $(Z)$ |
| $(H-)$ | Set of all non-carriers of the $(H+)$ HLA haplotype in $(Z)$ |
| $(M)$ | Set of all men in $(Z)$ |
| $(F)$ | Set of all women in $(Z)$ |
| $P(E)$ | Probability of an environmental exposure “sufficient to cause *MS*” in the set $(G)$, given the prevailing environmental conditions of the time |
| $P(MS\vert MZ_{MS})$ | Life-time probability of getting *MS* in an *MZ*-twin whose co-twin either has, or will develop, *MS* |
| $P(MS\vert IG_{MS})$ | Value of $P(MS\vert MZ_{MS})$, which has been adjusted to exclude the impact of the similar intrauterine and early post-natal environments of *MZ*-twins |

\* A complete presentation of the Model and more precise definitions of these terms are provided in *File S1*.

Nevertheless, each of these methods estimates only the prevalence of “diagnosed” MS, which may underestimate the prevalence of “pathological” MS in the population (*S1 File*). For example, several autopsy studies [34-37] have reported that “pathological” MS in patients is found in approximately 0.1% of individuals who were “undiagnosed” (i.e., they were either minimally symptomatic or asymptomatic) during life. In addition, at present, many of these asymptomatic individuals can now be identified (*in vivo*) using modern imaging methods [38]. As a result, the actual life time-risk of MS in the population is likely to be somewhat higher that this 0.3% estimate, and possibly by as much as 50-100% (*File S1*).

Also, within the general population (*Z*), we can define a subset of so-called “genetically-susceptible” individuals (*G*), such that every individual in this subset has a non-zero probability of developing MS (Table 1). All individuals who are not members of the (*G*) subset, therefore, are members of the so-called “non-susceptible” subset (*G*-).

In addition, we will define the term {*P*(*MS* | *MZ*_*MS*_)} to represent the life-time probability of developing MS for an individual from a monozygotic (*MZ*) twin-ship, given the fact that their identical co-twin either has or will develop MS (Table 1). This probability is estimated by the proband-wise concordance rate for *MZ* twins [39] and most epidemiological studies in northern populations (e.g., Table 2) report this proband-wise concordance rate to be approximately 25– 30% [40-46]. However, *MZ*-twins, in addition to sharing their identical genotypes (*IG*), also share similar intrauterine and early post-natal environments. Therefore, we define the term *P*(*MS* | *IG*_*MS*_) to represent the *MZ*-twin concordance rate, which has been adjusted to account for these environmental similarities. As developed in the *S1 File*, this adjustment can be made using the observed proband-wise concordance rates of siblings and fraternal twins. - i.e., siblings who share the same genetic relationship but are divergent in their intrauterine and early post-natal experiences [40,47-49].

**Table 2.** Canadian Population Data on *MZ*-Twin Concordance broken down by (*H+*)-haplotype and Gender-Status *

| ***HLA-DRB1\*15 Status*** | ***H+*** | ***H–*** | ***Totals*** |
| --- | --- | --- | --- |
| Concordant for MS (C) | 9 | 11 | 20 |
| Discordant for MS (D) | 31 | 42 | 73 |
| Totals | 40 | 53 | 93 |
| Pair-wise Concordance | 0.23 | 0.21 | 0.22 |
| Proband-wise Concordance | 0.31 | 0.29 | 0.30 |
| ***Gender Status*** | ***Women*** | ***Men*** | ***Totals*** |
| Concordant for MS (C) | 22 | 2 | 24 |
| Discordant for MS (D) | 66 | 43 | 109 |
| Totals | 88 | 45 | 133 |
| Pair-wise Concordance | 0.25 | 0.04 | 0.18 |
| Proband-wise Concordance | 0.34 | 0.07 | 0.25 |

***Other Canadian Data:***
\* Data from Willer et al. [40] and Orton et al. [68]
– the MZ-twins were drawn from the 19,938 MS-patients in the CCGPSMS database
Pair-wise concordance calculated as: $C/(C+D)$
Proband-wise concordance calculated as: $2C/(2C+D)$
– adjusted [39] for double ascertainment (13/24=54%)

We also define two other widely reported partitions of the general population. First, we designate those individuals who possess 1 or 2 copies of the Class II *HLA*-*DRB1*15:01~HLA*-*DQB1*06:02~a1* haplotype – i.e. the (*H+*) haplotype – as being members of the (*H+*) subset and those who possess 0 copies of this haplotype as being in the (*H-*) subset. Second, the partition consisting of women (*F*) and men (*M*) is considered. Finally, we consider *P*(*E*) the probability that (*G*)-subset members will experience environmental conditions sufficient to cause MS, given the prevailing environmental conditions of the time

The estimated values of, and the conditional relationship between, these different subsets is comprehensively developed in the *S1 File*. However, ten of the more notable conclusions are:

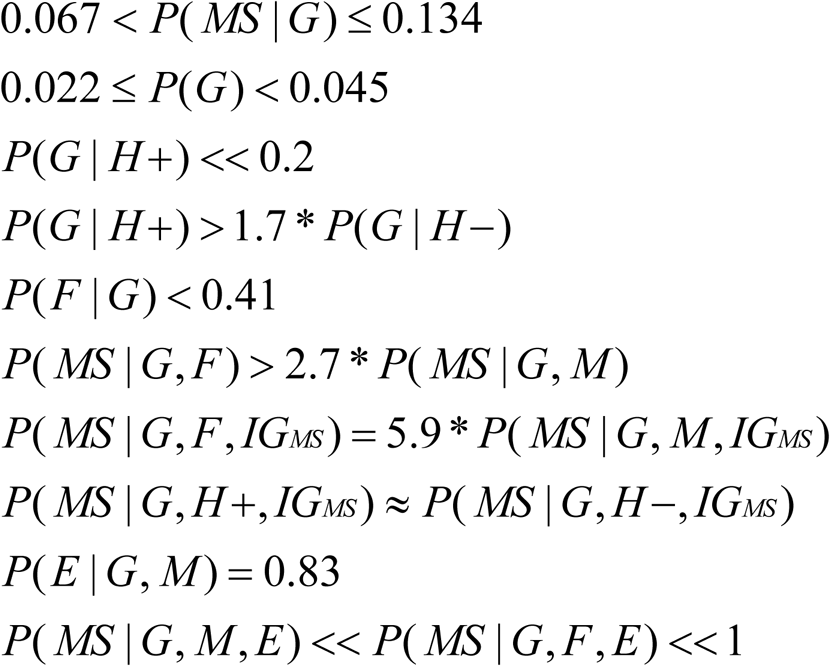

Each of these ten statements is unequivocal based on available epidemiologic data in Canada (*S1 File*). Notably, only 2.2–4.5% of the general population is even capable of getting MS – an estimate that is independently, and consistently, supported by epidemiological data from many populations throughout the northern hemisphere (*see File S1; Table S1*). This indicates that MS is fundamentally a genetic disorder. Moreover, the distribution of penetrance values within the general population (*Z*) is bimodal [50-54] with the large majority of individuals having no chance of getting MS and a small proportion – those individuals in the subset (*G*) – having a unique predisposition to MS (*File S1*).

Notable, also, is the fact that the vast majority of (*H+*)-carriers (<<20%) do not belong to the (*G*) subset. Therefore, at least with respect to the (*H+*) haplotype, “genetic-susceptibility” to MS must be the result of the combined effect of (*H+*) together with the effects of other genetic factors (*File S1*). By itself, the (*H+*) haplotype carries no MS-risk whatsoever. Finally, the impact of specific environmental events on the development of MS is also critical (*File S1*). Thus, in addition to being a genetic disease, MS is also an environmental disease. Both factors are necessary; neither alone is sufficient (*File S1*).

### Relative Risk Models in MS

There are two basic epidemiological models for the accumulation of disease-risk (*see File S2*), which have been widely utilized – the so-called additive and multiplicative risk models [55-59]. Nevertheless, actual epidemiological circumstances often don’t fall neatly into one model or the other. Indeed, the same basic probability model can be used to approximate either an additive or a multiplicative accumulation of risk [56]. The difference depends upon the definition of the term “no interaction” between the risk-factors [55-59]. In studies of the “genetic-susceptibility” to MS, multiplicative risk models have generally been utilized [60-62], although this choice may not be appropriate in all circumstances (*see File S2*).

Indeed, in practice, there are certain difficulties, which are encountered when trying to assess the appropriateness of either model. First, in a case-control studies (such as the WTCCC), because the incidence of the disease is not assessed (as it would be in a prospective cohort study), the actual *RRs* cannot be determined [63]. However, for a rare disease such as MS {e.g., where: *P*(*MS*)≈0.003}, the *ORs* and the *RRs* are almost identical [63] and, thus, can be used interchangeably.

Second, and more important, is the selection of an appropriate reference group for calculating the *RRs* (*File S2*). This choice will, necessarily, influence how well any observations fit into one or another of these risk models. As noted above and discussed further in the *S2 File*, the theoretical underpinnings for both the additive and multiplicative models arise from the same underlying probability assumptions [55-58], and are predicated on the notion that MS-risk for the different potential “risk-factors” is as great or greater than the “risk” in the reference group (*File S2*). This requires identifying the reference group with the lowest MS risk of any. Although the (*G–*) subset, by definition, has the lowest MS risk of any, this group (even if it could be identified) cannot be used as a reference because all *RRs* calculated with respect to it would, by definition, be either infinite or undefined. Indeed, the fact that, for MS, the (*G–*) subset is non-empty (*see File S1*), indicates that both of these “risk models” are, at a theoretical level, invalid for characterizing the accumulation of disease risk with an increasing number of disease-associated “risk” factors. Nevertheless, using a different reference group – i.e., one containing, at least, some members of the (*G*) subset - could be used to evaluate (approximately) whether either of these two models fits with the available data. In this circumstance, any subgroup consisting of only members of the (*G–*) subset would have an *RR* of zero. The subgroup with lowest risk of any that we identified in the WTCCC data was the (*AP\**) subset. Therefore, this group was used for the present analysis, despite the fact that a subgroup with an even smaller (non-zero) disease-risk seems likely to exist (*File S2*).

## Results

### The MHC

There were 146 CEHs in the HLA region that had 50 or more representations in the WTCCC dataset and these accounted for 48% of the total number (59,884) of CEHs present. Information on 45 of the CEHs, which were found in our previous study [24] to have some relationship to MS susceptibility, is provided in the *S2 File* (*Tables S2 & S3*). Of these, only the CEHs (*c1, c2, c3*, and *c5*) had a sufficient number of observations to assess the MS-risk of either homozygous combinations or combinations with each other. Therefore, these MS-associated CEHs were divided into five groups: 1) (*H+*) CEHs (i.e., containing the *HLA*-*DRB1*15:01~HLA*-*DQB1*06:02~a1* haplotype, *S2 File; Table S2*); 2) other increased risk or “extended risk” (*ER*) CEHs (*c23, c27, c34, c46, c68, c81, c85, c96*, and *c107*) CEHs as shown in *S2 File* (*Table S3*); 3) decreased risk or “all protective” (*AP*) CEHs (*c5, c15, c18, c24, c30, c32, c51*, and *c73*), as shown in *S2 File* (*Table S3*); 4) the “zero” group (*0*) consisting of all those CEHs which did not belong to the (*H+*), (*ER*), or (*AP*) groups; and 5) the (*c1*) CEH by itself. Each of these groups of CEHs seemed to be segregating independently and, in the control group, frequencies for each of the different combinations were, statistically, at their Hardy-Weinberg expectations.

The *MHC* allele *HLA-A*02:01* has been previously reported to be protective [64]. Although the association between *HLA-A*02:01*-positive status and MS was also found, in the WTCCC, to be “protective” relative to the (*0,0*) *MHC* genotype (*OR*=0.69; p<10^-29^), evidence from *Tables S2 & S3* (*S2 Fil*e) shows that any such association depends importantly upon the exact nature of the CEH on which this allele resides rather than upon the presence of the *HLA-A*02:01* allele itself. Thus, the grouping used here (i.e., the *AP* group) seems more appropriate than the use of this allele in isolation.

The subset of individuals who don’t carry any (*H+*), (*ER*), or (*AP*) CEHs at the *MHC* is referred to as the (*0,0*) *MH*C genotype. In Tables 2 & 3, all *ORs* are presented relative to this group. Each of the (*H+*) CEHs with 50 or more representations were significantly associated with MS-risk (*File S2*; *Table S2*), as were, collectively, (*H+*)-carrying CEHs with fewer than 50 representations in the WTCCC (Table 3). Moreover, also assessing, collectively, only those (*H+*)-carrying CEHs that a single representation in the WTCCC, the disease association is still statistically significant and of similar magnitude to other (*H+*)-carrying CEHs (i.e., *OR*=3.0; CI=2.7-3.4; p<10^-10^). Consequently, the (*H+*)-haplotype, by itself, seems to contribute to the disease susceptibility in an individual although, as shown in *File S2* (*Table S2*), the magnitude of this effect varies among different (*H+*)-carrying CEHs [25].

**Table 3.**
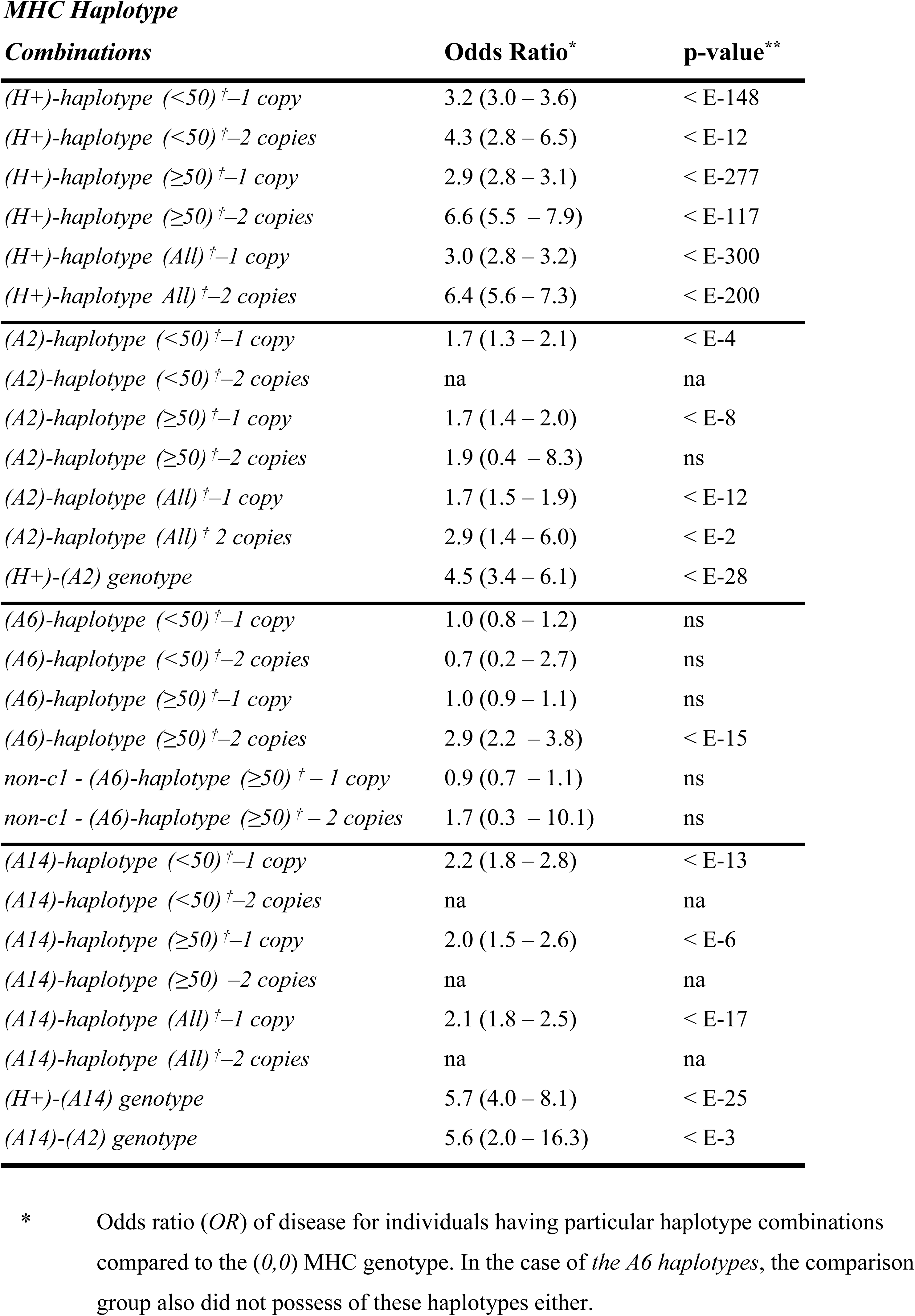
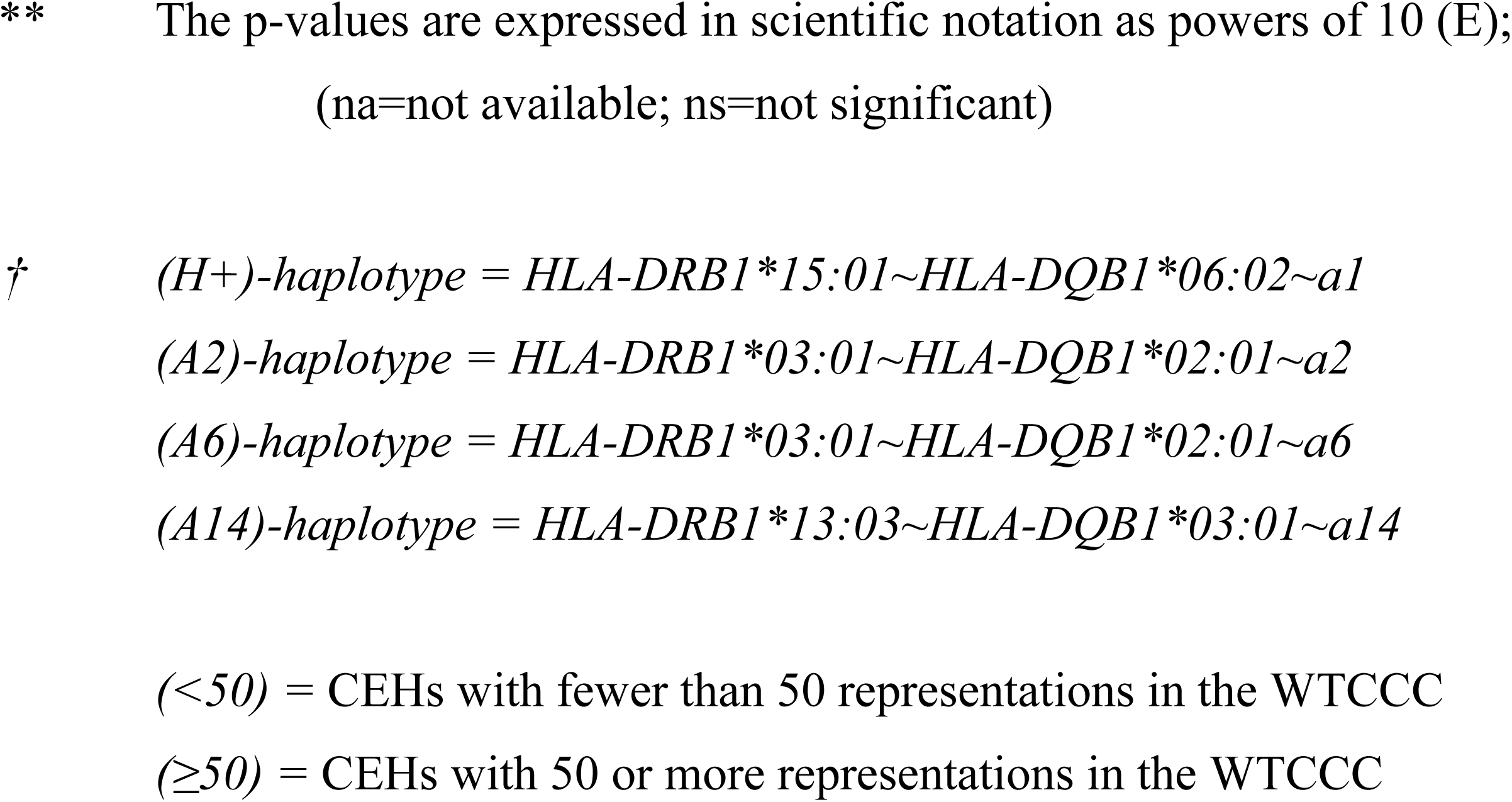
Impact of *CEH*s Frequency on Disease Association

| **Combinations** | **Odds Ratio\*** | **p-value\*\*** |
| --- | --- | --- |
| *(H+)*-haplotype (<50) <sup>†</sup>-1 copy | 3.2 (3.0 – 3.6) | < E-148 |
| *(H+)*-haplotype (<50) <sup>†</sup>-2 copies | 4.3 (2.8 – 6.5) | < E-12 |
| *(H+)*-haplotype (≥50) <sup>†</sup>-1 copy | 2.9 (2.8 – 3.1) | < E-277 |
| *(H+)*-haplotype (≥50) <sup>†</sup>-2 copies | 6.6 (5.5 – 7.9) | < E-117 |
| *(H+)*-haplotype (All) <sup>†</sup>-1 copy | 3.0 (2.8 – 3.2) | < E-300 |
| *(H+)*-haplotype (All) <sup>†</sup>-2 copies | 6.4 (5.6 – 7.3) | < E-200 |
| *(A2)*-haplotype (<50) <sup>†</sup>-1 copy | 1.7 (1.3 – 2.1) | < E-4 |
| *(A2)*-haplotype (<50) <sup>†</sup>-2 copies | na | na |
| *(A2)*-haplotype (≥50) <sup>†</sup>-1 copy | 1.7 (1.4 – 2.0) | < E-8 |
| *(A2)*-haplotype (≥50) <sup>†</sup>-2 copies | 1.9 (0.4 – 8.3) | ns |
| *(A2)*-haplotype (All) <sup>†</sup>-1 copy | 1.7 (1.5 – 1.9) | < E-12 |
| *(A2)*-haplotype (All) <sup>†</sup> 2 copies | 2.9 (1.4 – 6.0) | < E-2 |
| *(H+)*-(*A2*) genotype | 4.5 (3.4 – 6.1) | < E-28 |
| *(A6)*-haplotype (<50) <sup>†</sup>-1 copy | 1.0 (0.8 – 1.2) | ns |
| *(A6)*-haplotype (<50) <sup>†</sup>-2 copies | 0.7 (0.2 – 2.7) | ns |
| *(A6)*-haplotype (≥50) <sup>†</sup>-1 copy | 1.0 (0.9 – 1.1) | ns |
| *(A6)*-haplotype (≥50) <sup>†</sup>-2 copies | 2.9 (2.2 – 3.8) | < E-15 |
| *non-cl* - (*A6*)-haplotype (≥50) <sup>†</sup> - 1 copy | 0.9 (0.7 – 1.1) | ns |
| *non-cl* - (*A6*)-haplotype (≥50) <sup>†</sup> - 2 copies | 1.7 (0.3 – 10.1) | ns |
| *(A14)*-haplotype (<50) <sup>†</sup>-1 copy | 2.2 (1.8 – 2.8) | < E-13 |
| *(A14)*-haplotype (<50) <sup>†</sup>-2 copies | na | na |
| *(A14)*-haplotype (≥50) <sup>†</sup>-1 copy | 2.0 (1.5 – 2.6) | < E-6 |
| *(A14)*-haplotype (≥50) <sup>†</sup>-2 copies | na | na |
| *(A14)*-haplotype (All) <sup>†</sup>-1 copy | 2.1 (1.8 – 2.5) | < E-17 |
| *(A14)*-haplotype (All) <sup>†</sup>-2 copies | na | na |
| *(H+)*-(*A14*) genotype | 5.7 (4.0 – 8.1) | < E-25 |
| *(A14)*-(*A2*) genotype | 5.6 (2.0 – 16.3) | < E-3 |

\* Odds ratio (*OR*) of disease for individuals having particular haplotype combinations compared to the (0,0) MHC genotype. In the case of the *A6* haplotypes, the comparison group also did not possess of these haplotypes either.
\*\* The p-values are expressed in scientific notation as powers of 10 (E); (na=not available; ns=not significant)
† (H+)-haplotype = HLA-DRB1\*15:01~HLA-DQB1\*06:02~a1 (A2)-haplotype = HLA-DRB1\*03:01~HLA-DQB1\*02:01~a2 (A6)-haplotype = HLA-DRB1\*03:01~HLA-DQB1\*02:01~a6 (A14)-haplotype = HLA-DRB1\*13:03~HLA-DQB1\*03:01~a14
(<50) = CEHs with fewer than 50 representations in the WTCCC
(≥50) = CEHs with 50 or more representations in the WTCCC

In addition, as in Table 4 (*see legend*), we defined different “risk” CEH combinations as: “single copy risk” [1 copy of any (*H+*)-haplotype or any *ER* haplotype]; and: 2) “double copy risk” [2 copies of any (*H+*)-haplotype, (*c1*), or any *ER* haplotype, or any combination of {(*H+*) + *ER*}, {(*H+*) + (*c1*)}, or {*ER* + (*c1*)}]. The different “protective” CEH combinations were defined similarly as: 1) “single copy protective” [1 copy of an *AP* haplotype]; and: 2) “double copy protective” [2 copies of an *AP* haplotype]. Considering all of these “risk” CEH combinations (relative to the (*0,0*) MHC genotype), the (*H+*)-haplotypes accounted for 81% of the risk haplotypes in the control population and for approximately the same percentage of this risk in both men and women (80% and 82% respectively). Moreover, the likelihood of men in the control population possessing such a risk-CEH combination (26%) was approximately the same as the likelihood in women (27%). Similarly, the likelihood of men in the control population possessing an *AP* CEH (9%) was approximately the same as the likelihood in women (8%). Nevertheless, the “single copy risk” of MS for (*H+*)- and (*ER*)-haplotypes in women (OR=3.0; CI=2.8-3.2; p<10^-220^) was significantly greater (z=2.4; p=0.009) than the same risk in men (OR=2.6; CI=2.4-2,8; p<10^-96^). By contrast, the “double copy risk” of MS in women and men was about the same.

**Table 4.**
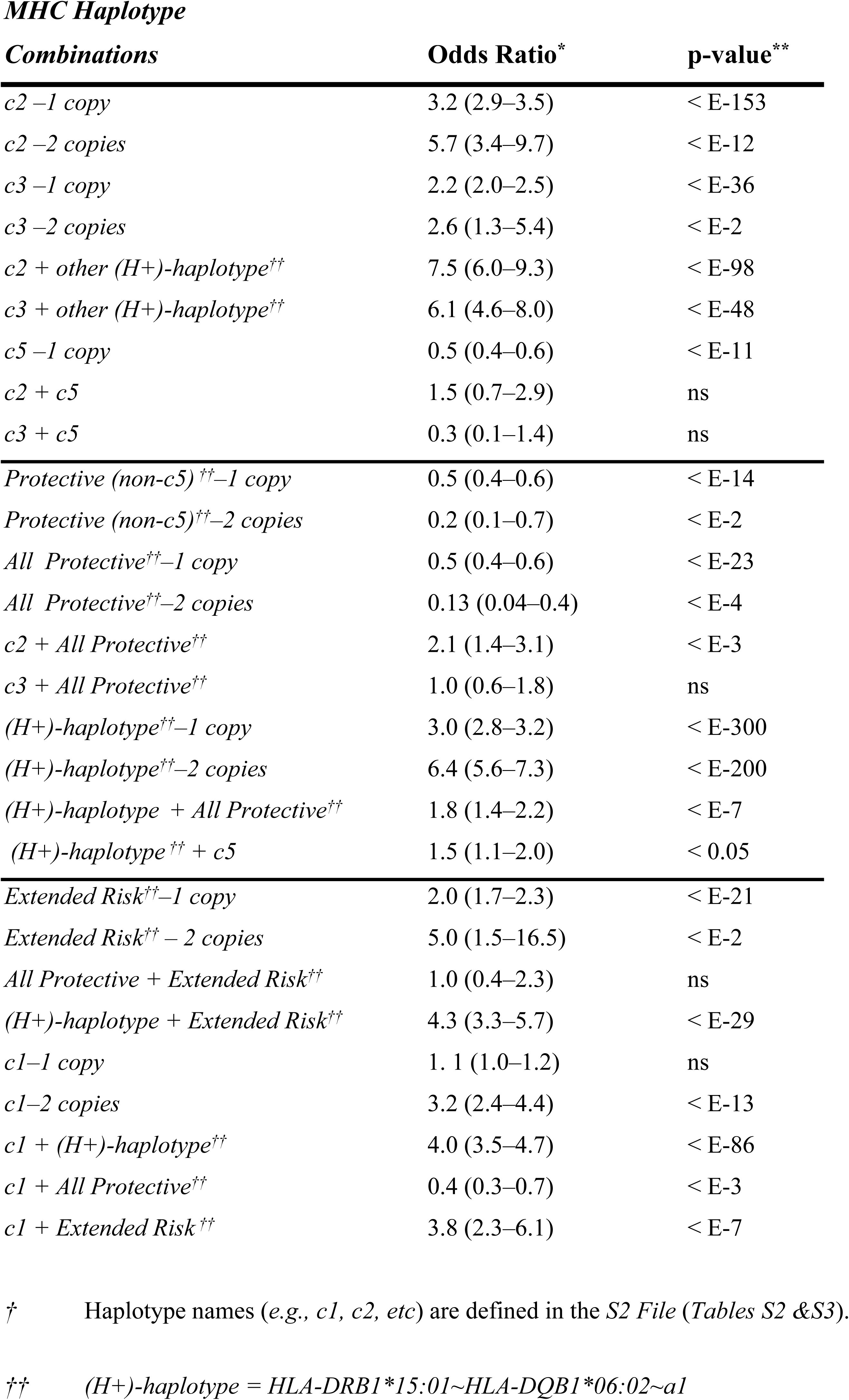
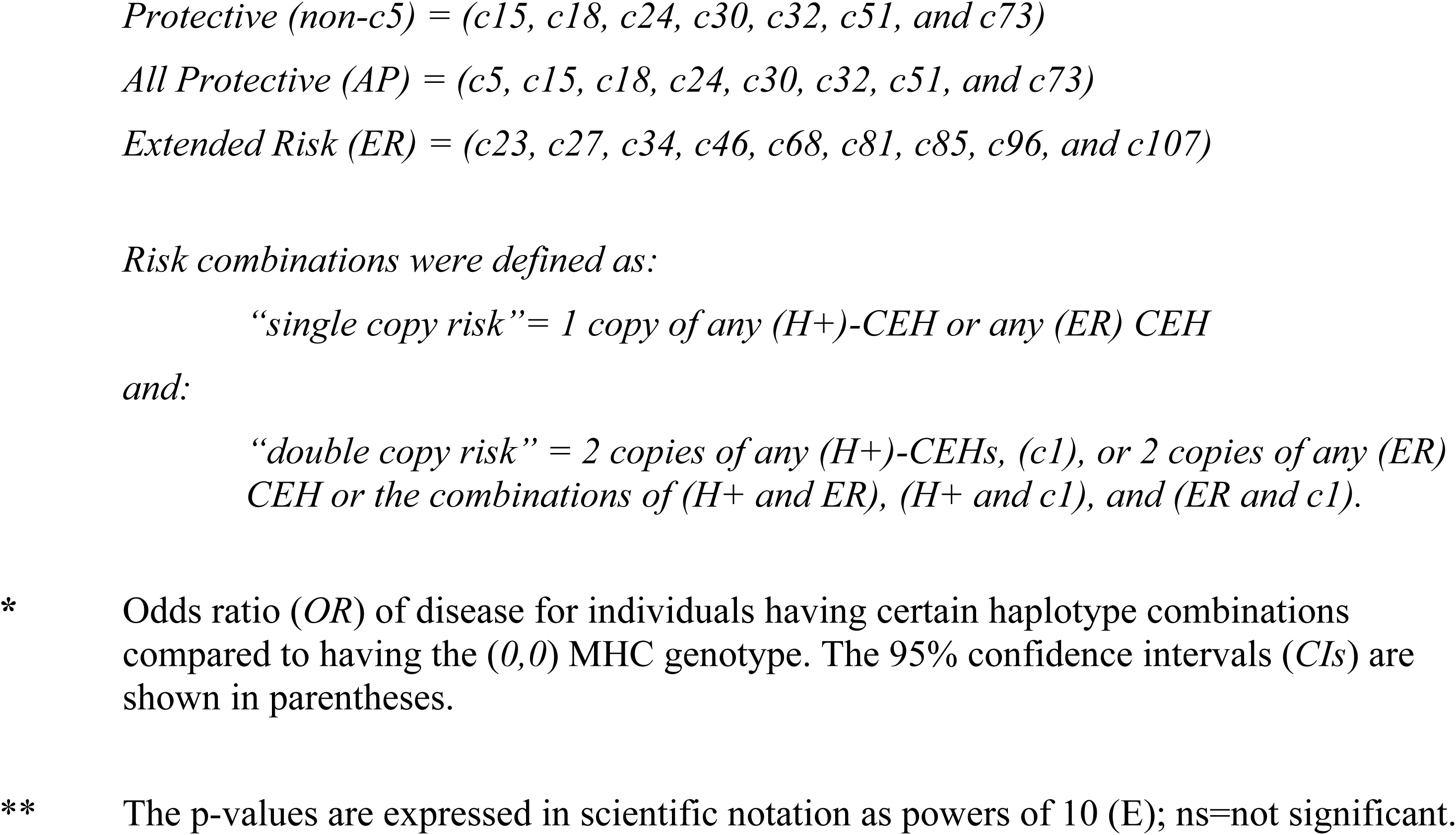
Impact on Phenotype of Combining MHC *CEH*s into a Genotype^†^

| **Combinations** | **Odds Ratio<sup>\*</sup>** | **p-value<sup>\*\*</sup>** |
| --- | --- | --- |
| *c2* –1 copy | 3.2 (2.9–3.5) | < E-153 |
| *c2* –2 copies | 5.7 (3.4–9.7) | < E-12 |
| *c3* –1 copy | 2.2 (2.0–2.5) | < E-36 |
| *c3* –2 copies | 2.6 (1.3–5.4) | < E-2 |
| *c2* + other (*H*+)–haplotype<sup>††</sup> | 7.5 (6.0–9.3) | < E-98 |
| *c3* + other (*H*+)–haplotype<sup>††</sup> | 6.1 (4.6–8.0) | < E-48 |
| *c5* –1 copy | 0.5 (0.4–0.6) | < E-11 |
| *c2* + *c5* | 1.5 (0.7–2.9) | ns |
| *c3* + *c5* | 0.3 (0.1–1.4) | ns |
| *Protective* (*non-c5*)<sup>††</sup>–1 copy | 0.5 (0.4–0.6) | < E-14 |
| *Protective* (*non-c5*)<sup>††</sup>–2 copies | 0.2 (0.1–0.7) | < E-2 |
| *All Protective*<sup>††</sup>–1 copy | 0.5 (0.4–0.6) | < E-23 |
| *All Protective*<sup>††</sup>–2 copies | 0.13 (0.04–0.4) | < E-4 |
| *c2* + *All Protective*<sup>††</sup> | 2.1 (1.4–3.1) | < E-3 |
| *c3* + *All Protective*<sup>††</sup> | 1.0 (0.6–1.8) | ns |
| (*H*+)–haplotype<sup>††</sup>–1 copy | 3.0 (2.8–3.2) | < E-300 |
| (*H*+)–haplotype<sup>††</sup>–2 copies | 6.4 (5.6–7.3) | < E-200 |
| (*H*+)–haplotype + *All Protective*<sup>††</sup> | 1.8 (1.4–2.2) | < E-7 |
| (*H*+)–haplotype<sup>††</sup> + *c5* | 1.5 (1.1–2.0) | < 0.05 |
| *Extended Risk*<sup>††</sup>–1 copy | 2.0 (1.7–2.3) | < E-21 |
| *Extended Risk*<sup>††</sup> – 2 copies | 5.0 (1.5–16.5) | < E-2 |
| *All Protective* + *Extended Risk*<sup>††</sup> | 1.0 (0.4–2.3) | ns |
| (*H*+)–haplotype + *Extended Risk*<sup>††</sup> | 4.3 (3.3–5.7) | < E-29 |
| *c1*–1 copy | 1.1 (1.0–1.2) | ns |
| *c1*–2 copies | 3.2 (2.4–4.4) | < E-13 |
| *c1* + (*H*+)–haplotype<sup>††</sup> | 4.0 (3.5–4.7) | < E-86 |
| *c1* + *All Protective*<sup>††</sup> | 0.4 (0.3–0.7) | < E-3 |
| *c1* + *Extended Risk*<sup>††</sup> | 3.8 (2.3–6.1) | < E-7 |

<sup>†</sup> Haplotype names (*e.g.*, *c1*, *c2*, *etc*) are defined in the *S2 File* (Tables *S2* & *S3*).
<sup>††</sup> (*H*+)–haplotype = *HLA-DRB1\*15:01~HLA-DQB1\*06:02~a1*
*Protective (non-c5) = (c15, c18, c24, c30, c32, c51, and c73)*
*All Protective (AP) = (c5, c15, c18, c24, c30, c32, c51, and c73)*
*Extended Risk (ER) = (c23, c27, c34, c46, c68, c81, c85, c96, and c107)*
*Risk combinations were defined as:*
*“single copy risk” = 1 copy of any (H+)-CEH or any (ER) CEH*
*and:*
*“double copy risk” = 2 copies of any (H+)-CEHs, (c1), or 2 copies of any (ER) CEH or the combinations of (H+ and ER), (H+ and c1), and (ER and c1).*
- \* Odds ratio (OR) of disease for individuals having certain haplotype combinations compared to having the (0,0) MHC genotype. The 95% confidence intervals (CIs) are shown in parentheses. - \*\* The p-values are expressed in scientific notation as powers of 10 (E); ns=not significant.

Similar to the (*H+*)-haplotypes, CEHs carrying the *HLA-*motifs:

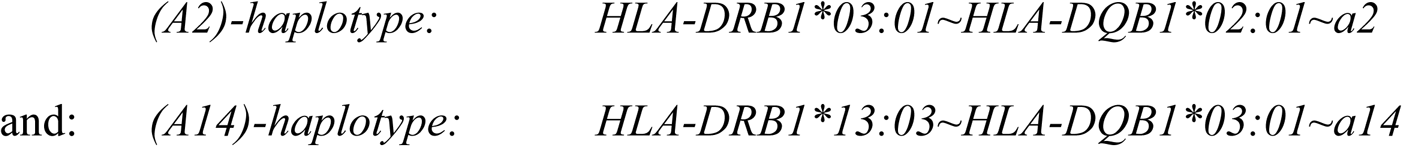

were associated with a disease risk that was similar regardless of the underlying frequency of the different CEHs (Tables 2 & 3). However, the same was not true for CEHs carrying the *HLA-* motif:

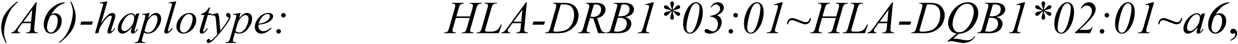

which seemed to vary quite widely in their disease association depending upon the exact CEH composition (Table 3; *S2 File; Table S3*).

The impact on the phenotype of an individual in response to combining two CEHs into a single genotype is shown in Table 4. For example, as has been well described previously [11,15-22], combining two copies of the (*H+*)-haplotype in to a single genotype markedly and significantly increases the disease association (Table 4; Fig. 1). Nevertheless, not all *(H+*)-carrying haplotypes have the same disease association [26]. For example, the *OR* for single copy carriers of the (*c2*) CEH is significantly greater (*z=*3.4–4.8; *p=*10^-3^– 10^-6^) than the *OR* for either single or double-copy carriers of the (*c3*) CEH.

**Figure 1.**
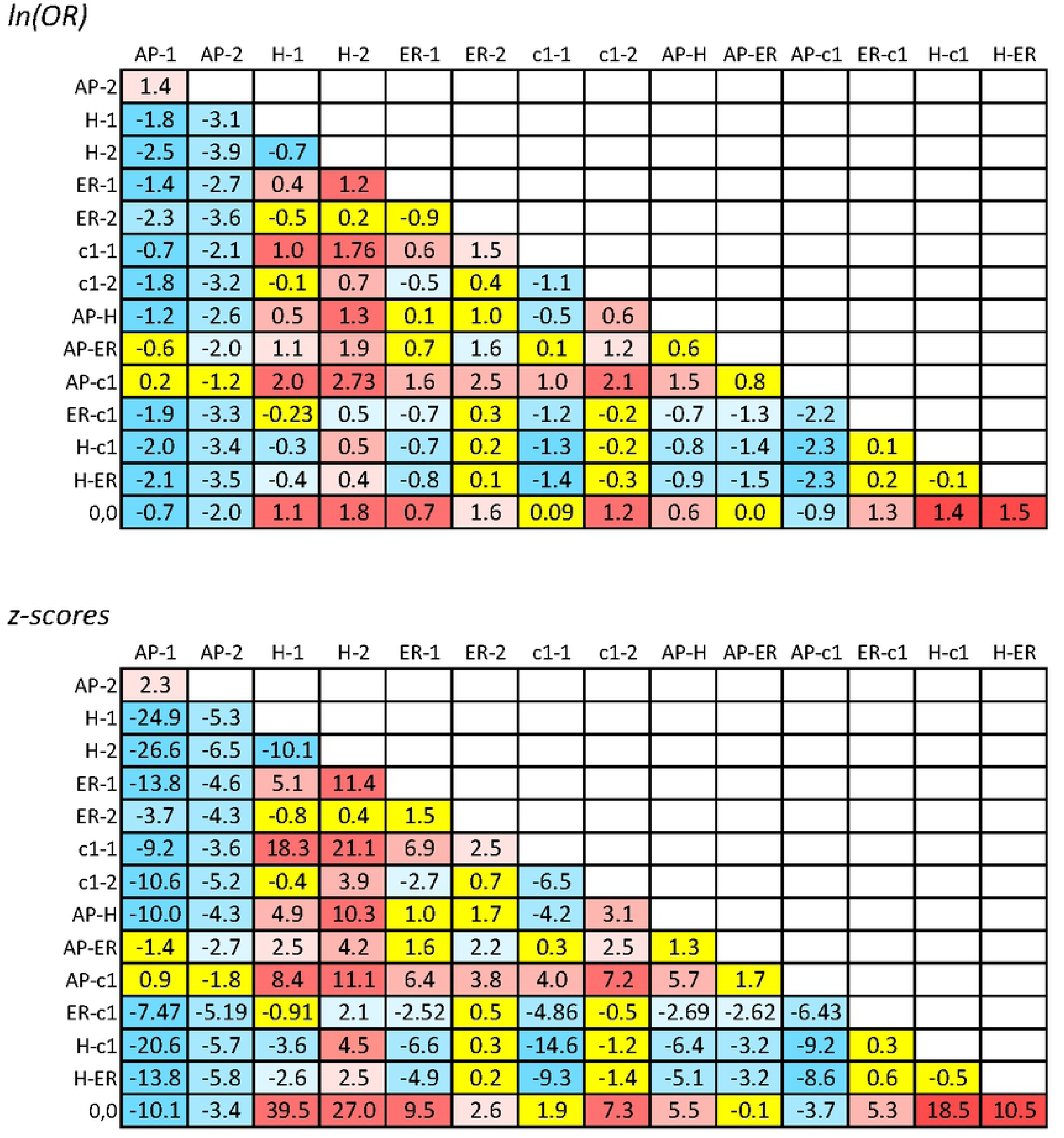
Lower triangular plot of the natural logarithm of odds ratios (*ORs*) and z-scores for the difference in disease risk for different two-*MHC* genotype combinations. To maximize statistical power, prior to calculating the comparative values for *ln*(*OR*), all *ORs* and standard deviations for each genotype were estimated relative to the *MHC* genotype (*0,0*) and then the combination compared to each other both as a ratio and as a z-score – see *Methods* and *Introduction*. Only genotypes with combinations of haplotypes with (*H+*), “extended risk” (*ER*), “all protective (*AP*), (*c1*), and (*0*) are shown. These genotypes are listed on both the *x-axis* (as columns) and *y-axis* (as rows) and the *ORs* and z-scores for each two-genotype comparison are represented as numbers at the points of intersection of the column and row for any two genotypes. Comparisons with an absolute z-score < 2.0, are shaded in yellow; comparisons with an absolute z-score = 2.0–3.0 are shaded in pale blue or pale red; comparisons with an absolute z-score = 3.1–7.0 are shaded in blue or red; comparisons with an absolute z-score > 7.0 are shaded in dark blue or dark red. Positive numbers (red shades) indicate that the genotype in the column has a greater *OR* than the genotype in the row. Conversely, negative numbers (blue shades) indicate that the genotype in the column has a smaller *OR* than the genotype in the row. For example, the genotype (*AP-2*) has a smaller *OR* than the genotype (*AP-1*). Similarly, the genotype (*H-1*) has a smaller *<u>OR</u>* than the combination (*H-2*). *ORs* given as (0.0) indicate that (*OR*<0.05). For no genotype were there zero MS cases observed. The following designations are use to indicate the different genotype configurations:

*AP1 = 1 copy of “All Protective” (AP) + 1 copy of (0)*
*AP2 = 2 copies of “All Protective” (AP)*
*H-1 = 1 copy of (H+) + 1 copy of (0)*
*H-2 = 2 copies of (H+)*
*ER-1 =1 copy of “Extended Risk” (ER) + 1 copy of (0)*
*ER-2 = 2 copies of “Extended Risk” (ER)*
*c1-1 = 1 copy of the c1 CEH + 1 copy of (0)*
*c1-2 = 2 copies of the c1 CEH*
*AP-ER = 1 copy of “AP” + 1 copy of “ER”*
*AP-H = 1 copy of “AP” + 1 copy of (H+)*
*AP-c1 = 1 copy of “AP” + 1 copy of the c1 CEH*
*ER-c1 = 1 copy of “ER” + 1 copy of the c1 CEH*
*H-c1 = 1 copy of (H+) + 1 copy of the c1 CEH*
*H-ER = 1 copy of (H+) + 1 copy of “ER*
*0,0 = 2 copies of (0)*

Similarly, considering the *AP* group of CEHs (Table 4; Fig. 1), we found a significant dose-dependent response such that possessing 2 copies of an *AP* CEH is significantly more “protective” than possessing only a single copy and, in addition, the magnitude of these “protective” effects is similar to the disease-risk produced by *(H+*)-haplotypes (Table 4; Fig. 1). Moreover, having an *AP* CEH, or even just the (*c5*) CEH, significantly and substantially mitigates (z=2.1–5.2; p=0.02–10^-7^) the disease risk produced by single copies of (*c2*), (*c3*), or, more generally, any (*H+*)-haplotype (Table 4; Fig. 1). A single copy of an “extended risk” CEH adds to the risk of a single copy of (*c2*), (*c3*), or any (*H+*)-haplotype, although it adds significantly less (z=2.5; p=0.006) than does a 2^nd^ copy of an (*H+*)-haplotype (Table 4; Fig 1). And, finally, the (*c1*) CEH acts in a recessive manner with little, if any, disease risk produced by a single copy (Table 4). Nevertheless, (and by contrast) a single copy of the (*c1*) haplotype adds significantly (z=2.5–6.0; p=0.006–10^-9^) to the disease risk produced by single copies of (*c2*), (*c3*), or, more generally, of any (*H+*)-haplotype (Table 4; Fig. 1).

Figure 2 shows the impact of replacing one MHC haplotype with another in different genotypic contexts. For example, replacing an (*0*)-haplotype with an (*H+*)-haplotype has a significantly greater impact when the companion is an (*0*)-haplotype compared to when the companion is an (*H+*)-haplotype (Fig. 2). Thus, comparing the (*0,H+*) genotype with the (*0,0*) genotype had an odds ratio of: (*OR*_*1*_*=3.0*) whereas, comparing the (*0,H+*) genotype with the (*H+,H+*) genotype had an odds ratio of: (*OR*_*2*_*=2.1*). These two *ORs* were significantly different from each other (*z=4.7*) and had a ratio of: *OR*_*1*_ */ OR*_*2*_*=1.4*; and: *ln(1.4*)=*0.4*.

**Figure 2.**
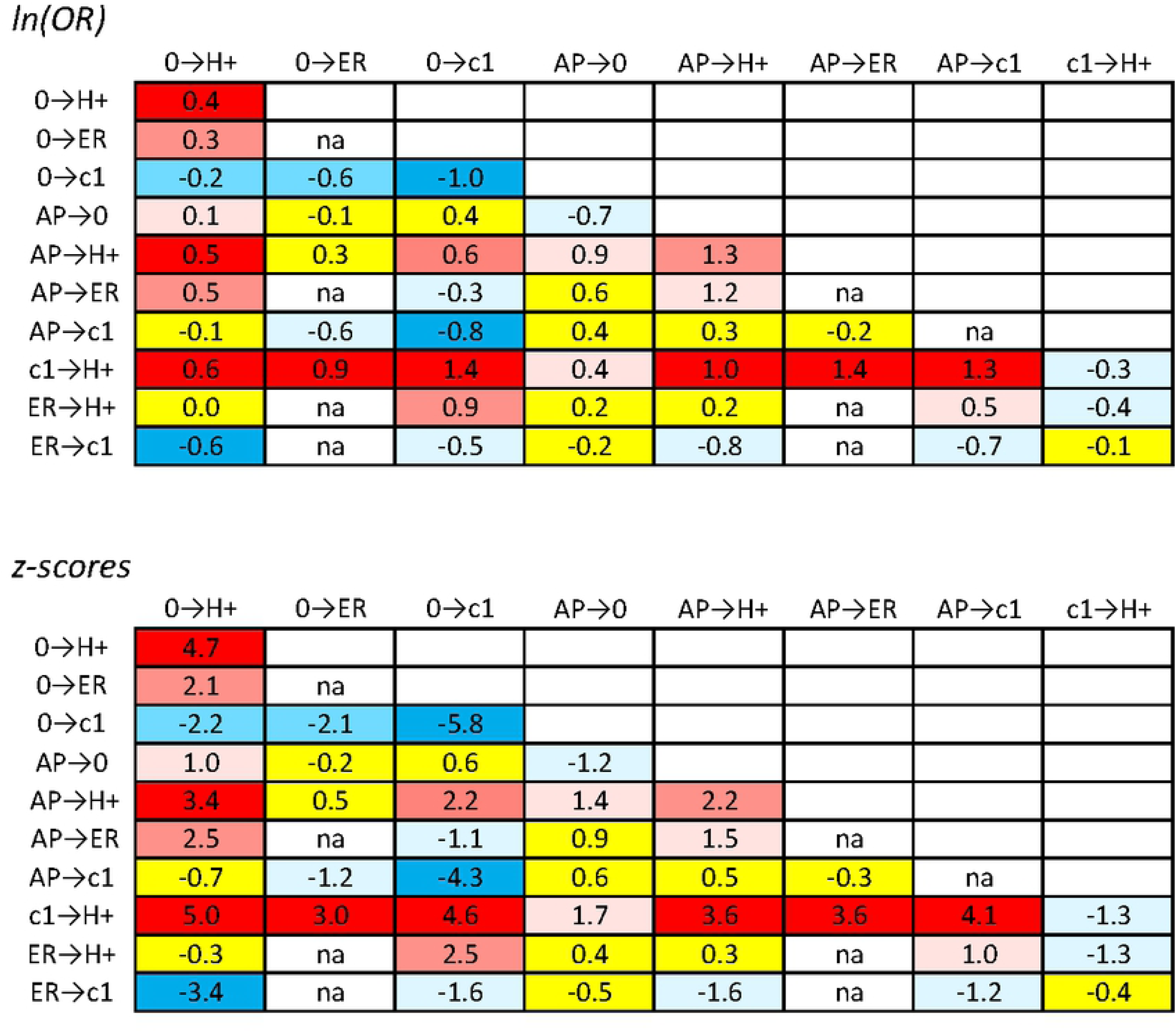
Lower triangular plots of the natural logarithm of the odds ratios (*ORs*) and z-scores for the different transitions from one MHC haplotype to another in different genotypic contexts. For example, the point of intersection for (0 → *H* +) and (*c*1 → *H* +) represents the ratio of:

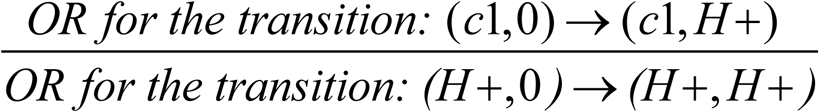

or equivalently:

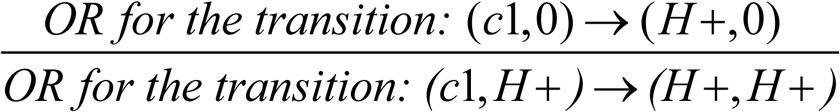 Only transitions for (*H+*), “extended risk” (*ER*), “all protective (*AP*), (*c1*), and (*0*) haplotypes are shown. These transitions are indicated on both the *x-axis* (as columns) and *y-axis* (as rows) and values for *ln*(*OR*) and the z-scores for each transition comparison are represented as numbers at the points of intersection of the column and row for any two genotypes. The designation “*na*” indicates data “not available”. Comparisons with an absolute z-score >3.0, are shaded either dark blue (negative) or dark red (positive); comparisons with an absolute z-score = 2.0–3.0 are shaded either light blue (negative) or light red (positive); comparisons with an absolute z-score = 1.0–2.0 are shaded either pale blue (negative) or pale red (positive) yellow; comparisons with absolute z-scores <1.0 are shaded in yellow; Positive numbers indicate that *OR* for the transition (indicated by the column) is greater for the 1^st^ genotypic configuration (indicated by the row) than it is for the 2^nd^. Conversely, a negative number indicates that the transition *OR* for the 2^nd^ genotypic configuration is greater than the 1^st^. For example, the number (3.4) in the 1^st^ column, 5^th^ row of the z-score table, indicates that the *OR* for transition from genotype (*AP,0*) to (*AP,H+*) is significantly greater that the *OR* for the transition from (*H+,0*) to (*H+,H+*). Similarly, the number (–2.2) in the 1^st^ column, 3^rd^ row of the z-score table, indicates that the *OR* for transition from genotype (*0,0*) to (*0,H+*) is significantly less that the *OR* for the transition from (*c1,0*) to (*c1,H+*). Because of symmetry, the *OR* for comparing the transition from genotype (*AP,0*) to (*AP,H+*) with the transition from genotype (*H+,0*) to (*H+,H+*) is mathematically equivalent to the *OR* for comparing the transition from genotype (*AP,0*) to (*H+,0*) with the transition from genotype (*AP,H+*) to (*H+,H+*). Therefore, the interpretation for the meaning of the rows and columns is completely interchangeable (although the implication of positive and negative numbers remains unchanged).

By contrast, replacing an (*0*)-haplotype with an (*H+*)-haplotype has a significantly smaller impact when the companion is an (*0*)-haplotype compared to when the companion is an (*c1*)-haplotype (Fig. 2). Thus, comparing the (*0,H+*) genotype with the (*0,0*) genotype had an odds ratio of: (*OR=3.0*) whereas, comparing the (*0,c1*) genotype with the (*H+,c1*) genotype had an odds ratio of: (*OR=3.7*). These two *ORs* were significantly different from each other (*z=-2.2*) and had a ratio of: *OR*_*1*_ / *OR*_*2*_*=0.8*; and: *ln(0.8*)=-*0.2*.

As can be appreciated from the figure, the impact of replacing one haplotype with another often depends considerably (and significantly) upon the exact nature of the companion haplotype, which, together with the haplotype being replaced, constitutes the *MHC* genotype (Fig. 2). This reflects the multiple haplotype-haplotype interactions that exist within the *MHC*. Indeed, if no such interactions were present, each of the comparisons provided in the figure would have an *OR* of ~1.0 - i.e., *ln*(*OR*)=0 - and would be shaded in yellow (Fig 2).

### The Non-*MHC* Loci

In the WTCCC data set, and as described previously [24], the (*d1*) haplotypes are 11-SNP haplotypes in Region #234, which consist of 185 different SNP combinations and, of which, 1,243 (2%) are the “risk” haplotype (01100000100); the (*d2*) haplotypes are 3-SNP haplotypes in Region #734, which consist of 7 different SNP combinations and, of which, 14,091 (23%) were the “risk” haplotype (111); and the (*d3*) haplotypes are 15-SNP haplotypes in Regions #814, #818 and #822, which consist of 210 different SNP combinations and, of which, 24,709 (41%) are the “risk” haplotype (000010000000000). The *ORs* for the various combinations of the non-*MHC* loci are shown in Table 5. The increase in disease susceptibility that results from combining susceptibility genotypes at these three non-*MHC* loci with MHC genotypes is quite different for the different MHC configurations (Fig 3). Thus, for example, the different combinations of these non-*MHC* “risk” haplotypes consistently increased the risk for (*0,H+*), (*H+,H+*), (*0,c1*), and (*H+,c1*) “risk” genotypes (Fig.3). By contrast, for other “risk” genotypes such as (*AP,H+*) and (*ER,H+*) and for “protective” genotypes such as (*AP,0*) and (*AP,c1*), these other these non-*MHC* “risk” haplotypes seemed to contribute essentially nothing to the final risk (Fig. 3).

**Table 5.**
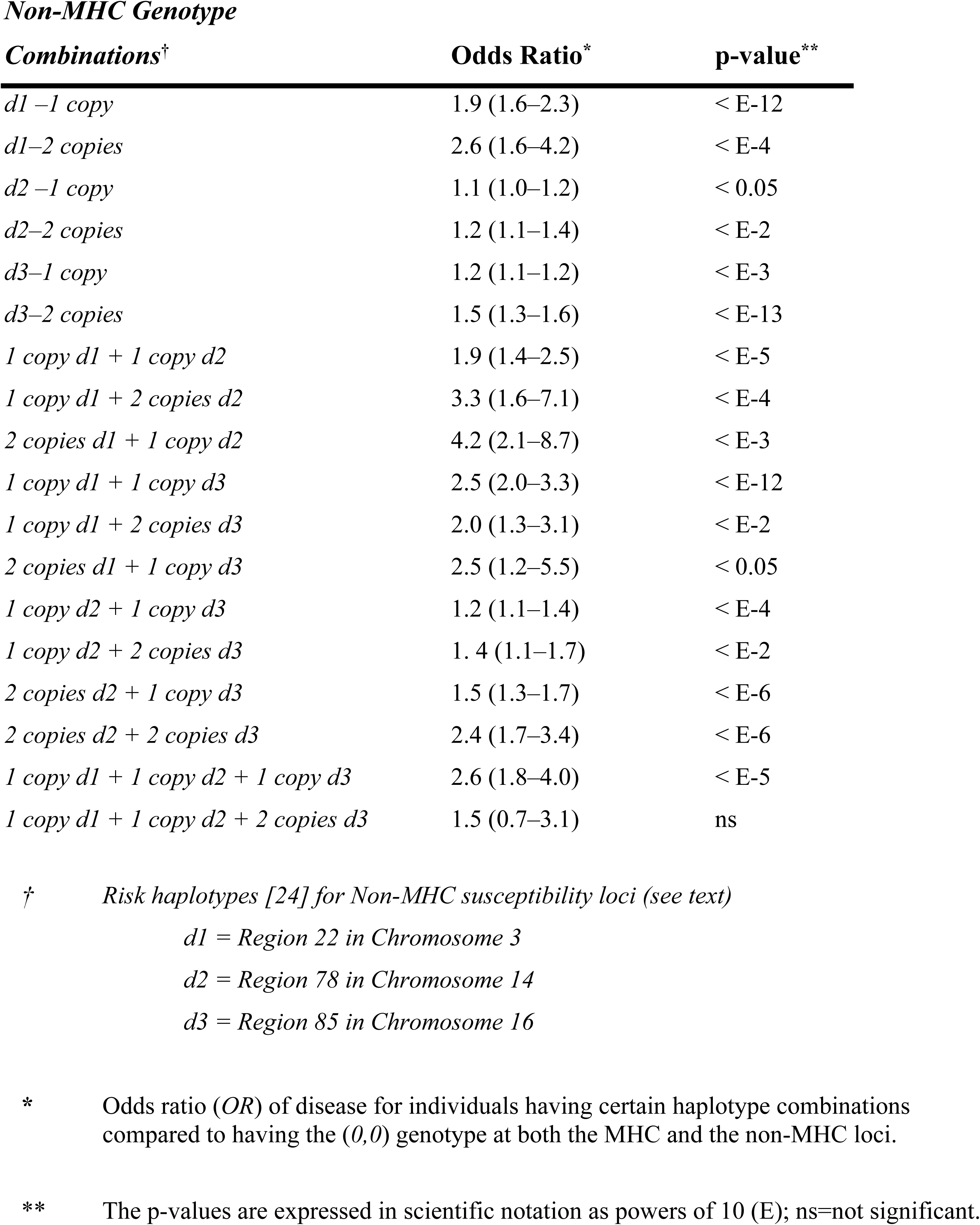
Impact on Phenotype of Combining Non-MHC Genotypes^†^

| **Combinations<sup>†</sup>** | **Odds Ratio<sup>\*</sup>** | **p-value<sup>\*\*</sup>** |
| --- | --- | --- |
| *d1 –1 copy* | 1.9 (1.6–2.3) | < E-12 |
| *d1–2 copies* | 2.6 (1.6–4.2) | < E-4 |
| *d2 –1 copy* | 1.1 (1.0–1.2) | < 0.05 |
| *d2–2 copies* | 1.2 (1.1–1.4) | < E-2 |
| *d3–1 copy* | 1.2 (1.1–1.2) | < E-3 |
| *d3–2 copies* | 1.5 (1.3–1.6) | < E-13 |
| *1 copy d1 + 1 copy d2* | 1.9 (1.4–2.5) | < E-5 |
| *1 copy d1 + 2 copies d2* | 3.3 (1.6–7.1) | < E-4 |
| *2 copies d1 + 1 copy d2* | 4.2 (2.1–8.7) | < E-3 |
| *1 copy d1 + 1 copy d3* | 2.5 (2.0–3.3) | < E-12 |
| *1 copy d1 + 2 copies d3* | 2.0 (1.3–3.1) | < E-2 |
| *2 copies d1 + 1 copy d3* | 2.5 (1.2–5.5) | < 0.05 |
| *1 copy d2 + 1 copy d3* | 1.2 (1.1–1.4) | < E-4 |
| *1 copy d2 + 2 copies d3* | 1.4 (1.1–1.7) | < E-2 |
| *2 copies d2 + 1 copy d3* | 1.5 (1.3–1.7) | < E-6 |
| *2 copies d2 + 2 copies d3* | 2.4 (1.7–3.4) | < E-6 |
| *1 copy d1 + 1 copy d2 + 1 copy d3* | 2.6 (1.8–4.0) | < E-5 |
| *1 copy d1 + 1 copy d2 + 2 copies d3* | 1.5 (0.7–3.1) | ns |

<sup>†</sup> Risk haplotypes [24] for Non-MHC susceptibility loci (see text)
*d1* = Region 22 in Chromosome 3
*d2* = Region 78 in Chromosome 14
*d3* = Region 85 in Chromosome 16
<sup>\*</sup> Odds ratio (OR) of disease for individuals having certain haplotype combinations compared to having the (0,0) genotype at both the MHC and the non-MHC loci.
<sup>\*\*</sup> The p-values are expressed in scientific notation as powers of 10 (E); ns=not significant.

**Figure 3.**
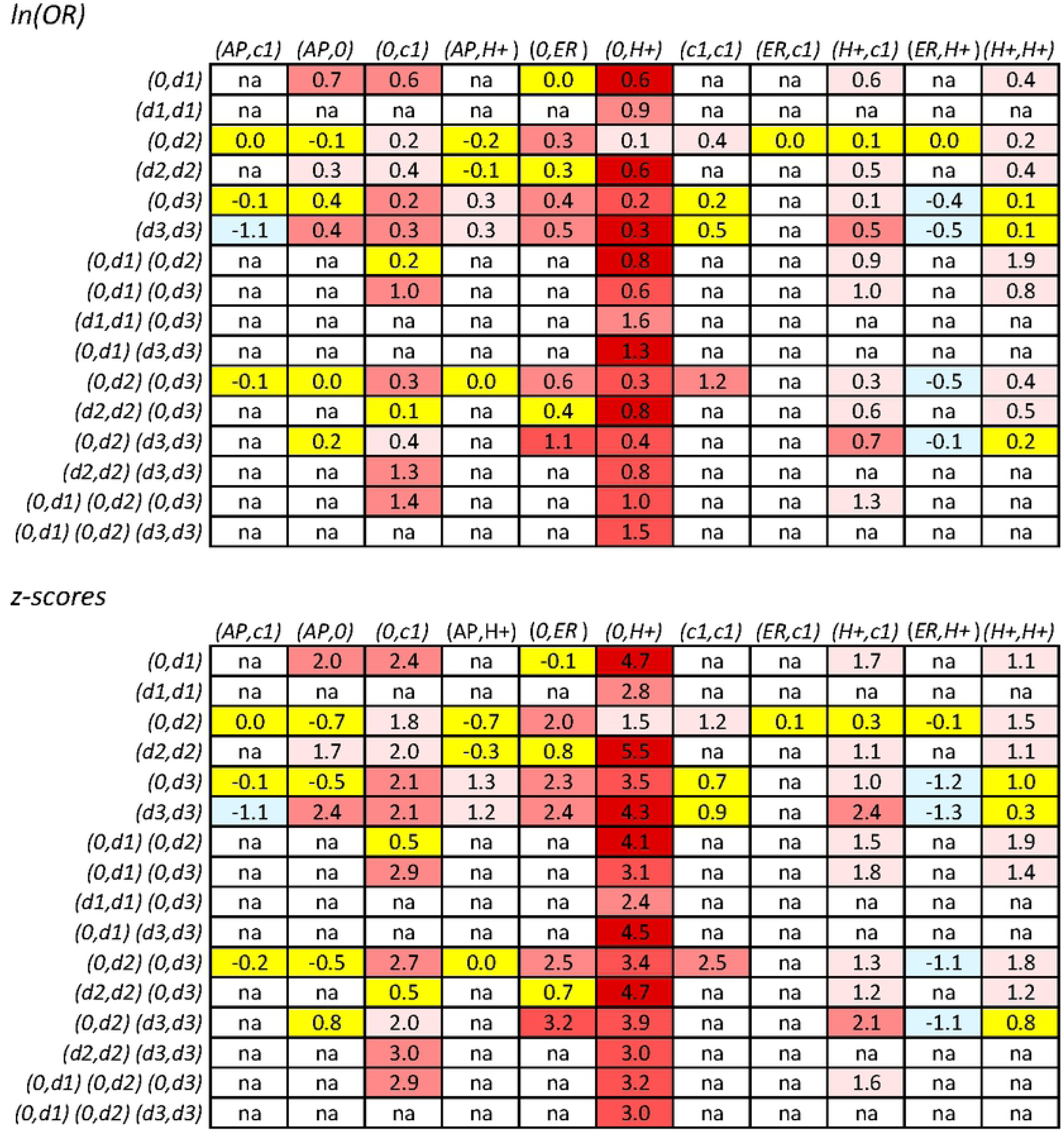
Natural logarithm of the odds ratios (*ORs*) and z-scores for the different combinations of *MHC* and non-*MHC* genotypes. All *ORs* were calculated relative to a group consisting of the same *MHC* genotype and with the genotypes (*0,0*) at all non-MHC loci involved in the comparison – see text. The *MHC* genotypes, in order of increasing disease-risk (Table 4), are presented on the *x-axis* (as columns) and the genotypes at non-*MHC* loci are presented on the *y-axis* (as rows). The values for *ln*(*OR*) and the *z-scores* for each comparison are represented as numbers at the points of intersection of the column and row for any two haplotypes. Comparisons with a z-score (|*z|<1*) are shaded in yellow; comparisons with a z-score (*1≥|z|≤2*) are shaded in either pale blue (negative) or pale red (positive); comparisons with a z-score (*2≥z≤3*) are shaded in light red; comparisons with a z-score (*3≥z≤4*) are shaded in red; comparisons with a z-score (*z≥4*) are maroon. Specific combinations having marginal totals of less than15 representations in the WTCCC are indicated by (*na*). Of the 83 observations presented, only 12% had a marginal total of less than 25 and 13% had a marginal total from 25 to less than 50.

### Additive vs. Multiplicative Risk

Combinations of the 3 non-*MHC* susceptibility regions, together with different genotypes at the MHC are presented in Figs. 4–7. In each of these Figures, the *ORs* are those derived from a comparison with (*AP,AP*) *MHC* genotype individuals as the reference. In all cases, the disease risk conferred by each genotype at each locus is estimated directly from the WTCCC observations (*see Methods*). The expectations from the additive and multiplicative risk-models are then compared to the actual observations (Figs. 4–7). In almost all cases, the additive model fits better with the actual observations than does the multiplicative model, especially as more “risk” loci are included in the combinations (Figs. 4–7). Nevertheless, neither model fits perfectly. When considering only *MHC* “risk” genotypes, for combinations of *MHC* genotypes whose disease-risk exceeds that of the (*0,0*) *MHC* genotype, the actual disease-risk observed is, in general, greater than predicted by the additive model (Fig. 4). By contrast, considering also the other non-*MHC* “risk” genotypes, the observed disease-risk is generally less than predicted by the additive model (Fig 5). This effect is increased when more “risk” loci are included in the combinations (Figs. 6,7).

**Figure 4.**
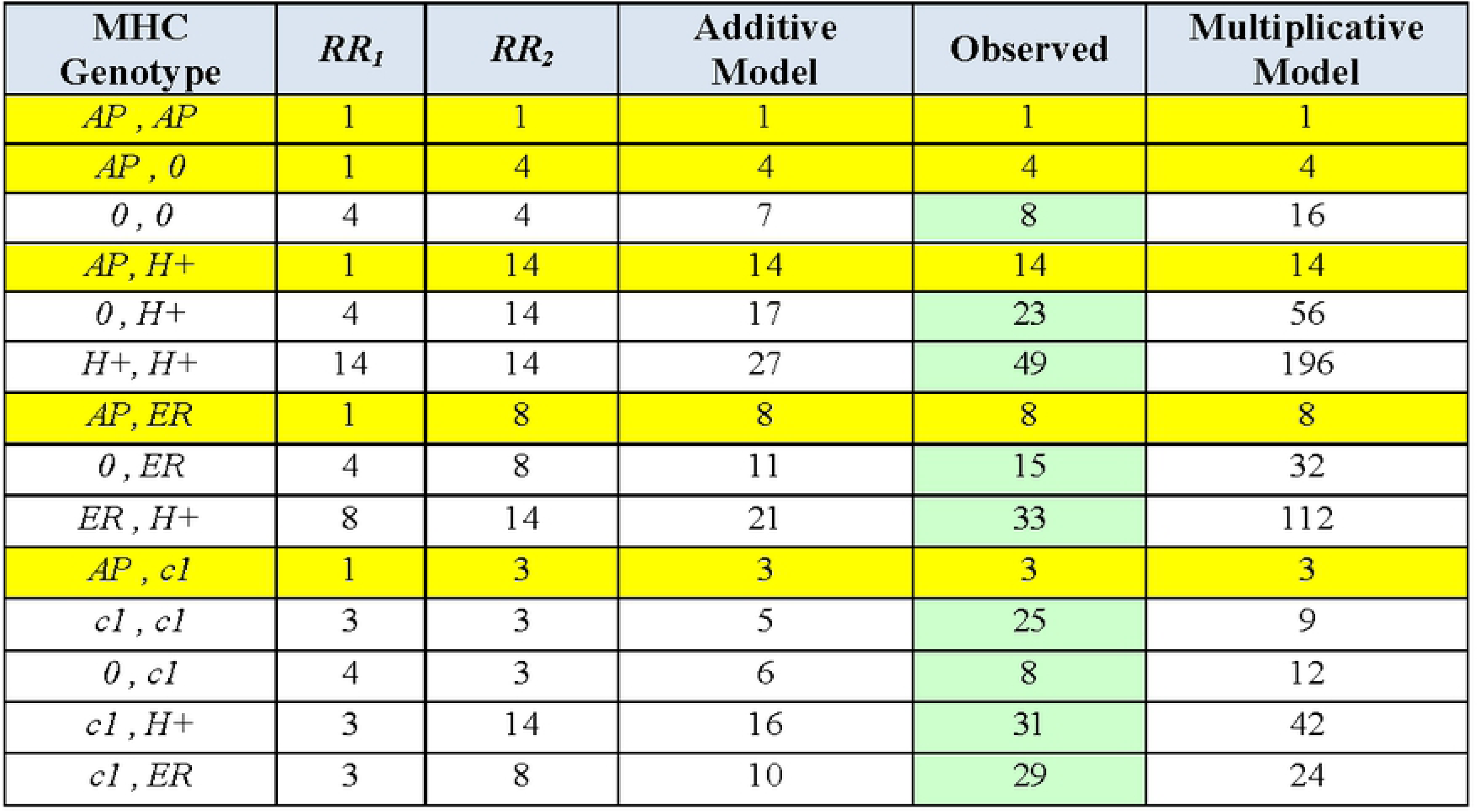
Conformity of the observed effect of combining different MHC haplotypes with an additive and a multiplicative model of combined risk. Yellow bands represent the definitional odds ratios (*ORs*) relative to a reference group consisting of the (*AP,AP*) or (*AP*^*\**^) genotype (i.e., as defined in the text: *R*_*b*_*=R*_*AP\**_*=*1). With the exception of (*c1*), which seems to behave in a unusual fashion, the combination of other risk alleles produced, in general, a risk in between the two models, albeit closer to that predicted by the additive model. All combinations had, at least, 50 representations in the WTCCC and the green shading indicates the *ORs* actually observed. Cells with yellow shading in the “Observed” column also represents the *ORs* actually observed. However, in these yellow-highlighted cases, the *ORs* were used to approximate the relative risks (*RRs*), which, in turn, were used to assess whether the genotypes that are not yellow-highlighted conformed to the additive and multiplicative models (*see Methods*).

**Figure 5.**
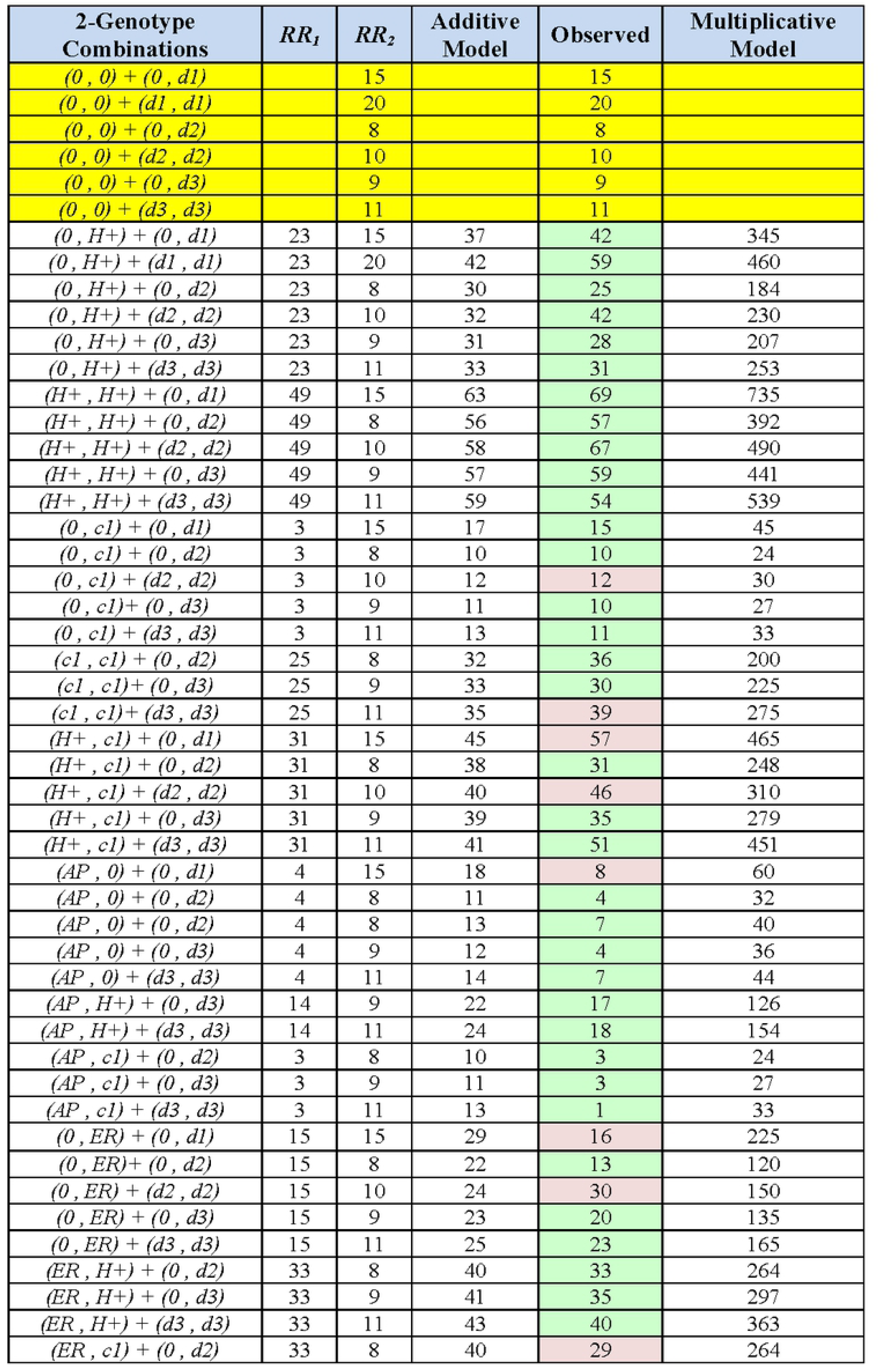
Conformity of the observed effect of combining different genotypes at the MHC and one susceptibility region with an additive and a multiplicative model of combined risk. The non-MHC susceptibility haplotypes are: (*d1*); (*d2*); and (*d3*) *–* see *Methods*. Yellow bands, as in Fig. 4, represent the definitional *ORs* for different non-MHC genotypes actually observed, but which have been re-referenced to a group with the (*AP,AP*) MHC genotype. The *ORs* for all MHC genotypes are also those actually observed (Fig 4). Only haplotype combinations with *≥*15 or more representations in the WTCCC are shown. Combinations with fewer than 50representations are shaded in pink; combinations with at least 50 representations are shaded in green.

**Figure 6.**
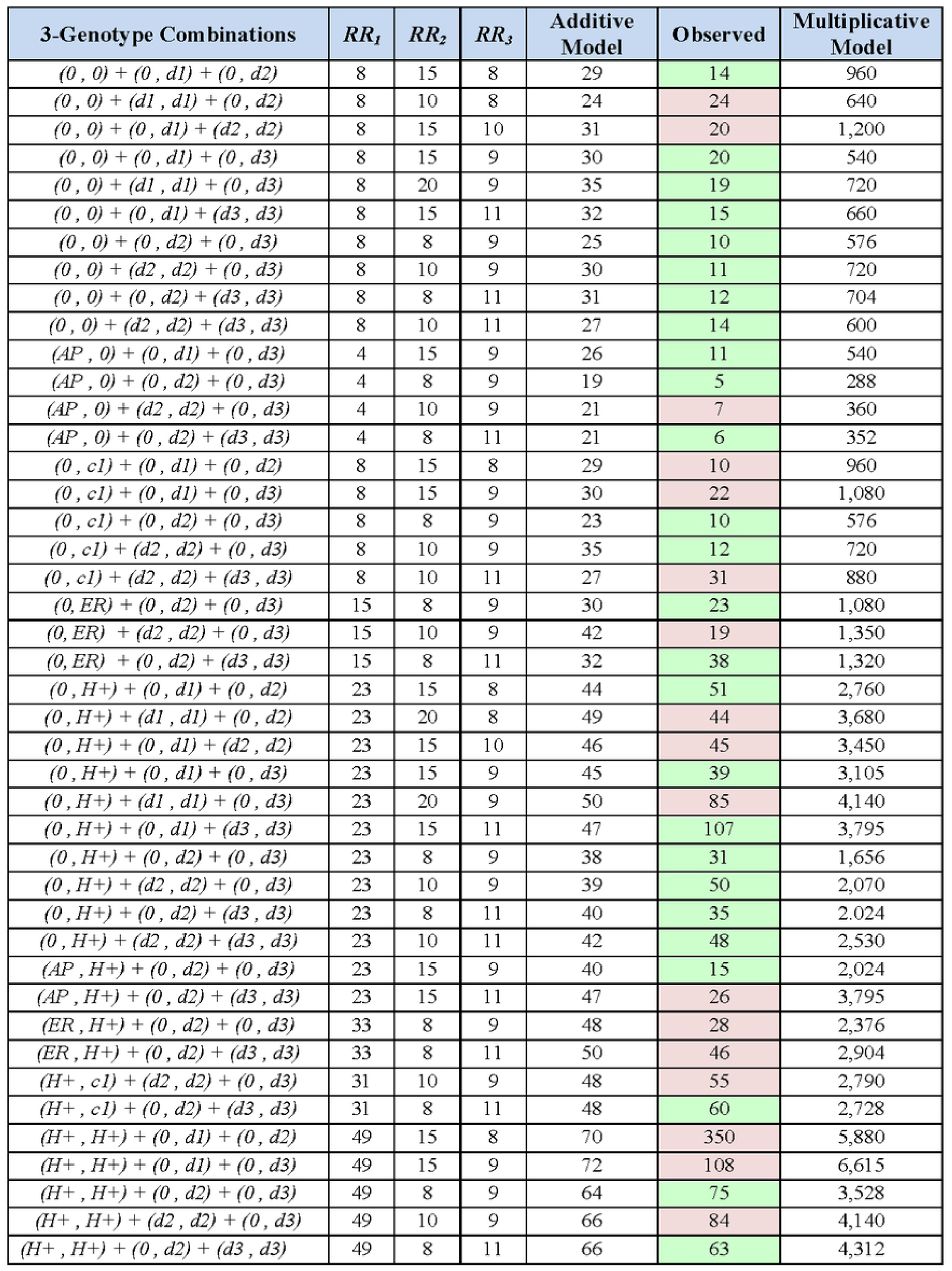
Conformity of the observed effect of combining different genotypes at the MHC and two susceptibility regions with an additive and a multiplicative model of combined risk. The non-MHC susceptibility haplotypes are: (*d1*); (*d2*); and (*d3*) *–* see *Methods*. The *ORs* listed are those actually observed (Figs 4 & 5). Only haplotype combinations with *≥*15 or more representations in the WTCCC are shown. Combinations with fewer than 50 representations are shaded in pink; combinations with at least 50 representations are shaded in green.

**Figure 7.**
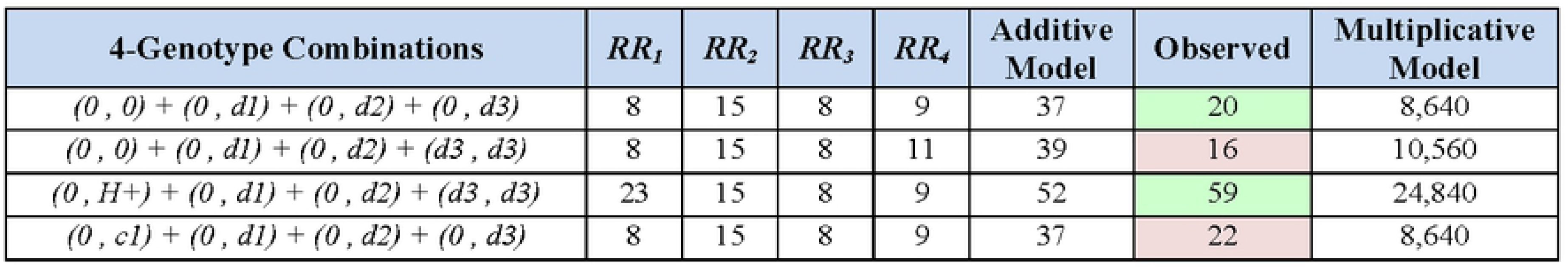
Conformity of the observed effect of combining different genotypes at the MHC and three susceptibility regions with an additive and a multiplicative model of combined risk. The non-MHC susceptibility haplotypes are: (*d1*); (*d2*); and (*d3*) *–* see *Methods*. The *ORs* listed are those actually observed (Figs 4 & 5). Only haplotype combinations with *≥*15 or more representations in the WTCCC are shown. Combinations with fewer than 50 representations are shaded in pink; combinations with at least 50 representations are shaded in green.

## Discussion

The present findings provide considerable insight to the underpinnings of “genetic-susceptibility” to MS and indicate that this susceptibility is complex. At the *MHC*, there are multiple different CEHs that contribute to susceptibility in different ways. When referenced to the *MHC* genotype (*0,0*), certain groups of CEHs seem to affect disease risk in a manner that either increase or decrease disease-risk when combined into a single genotype. For example, the combination of 2 “risk” CEHs (*H+* or *ER*) results in an increased disease risk compared to a single copy of a “risk” CEH alone (Tables 2 and 3, Fig. 1). Similarly, the combination to 2 “protective” CEHs (“protective” or “all protective”), results in a decreased disease risk compared to a single copy (Table 4; Fig. 1). Finally, combining a “risk” CEH together with a “protective” CEH results in an intermediate disease risk compared with having a single copy of either CEH-type alone (Table 4).

Nevertheless, there are exceptions to this general rule. Notably, when referenced to the (*0,0*) *MHC* genotype, a single copy of the (*c1*) CEH - the highest frequency CEH in both the WTCCC controls and other European populations [25,26] - is associated with a negligible, non-significant, disease-risk (Table 4; Fig. 1). By contrast, the disease-risk is substantially (and significantly) increased in the homozygous state (Table 4; Fig. 1). Such a pattern suggests that (*c1*) is acting in a recessive manner. Nevertheless, a single (*c1*) CEH increases disease risk when combined with “risk” CEHs but not with “protective” CEHs (Table 4). Thus, (*c1*) with an (*H+*) or an (*ER*) CEH resulted in a significantly increased disease risk compared to each CEH alone (Fig. 1). By contrast, the combination of (*c1*) and an (*AP*) CEH neither enhances nor to mitigates the effect of the “protective” CEH by itself (Fig.1).

Our findings have certain implications with respect to the appropriateness of the additive and multiplicative causal models for the accumulation of genetic risk. For appropriate *OR* or *RR* comparisons, calculations, need to be made using the lowest, non-zero, MS-risk as a reference group. In the WTCCC, the (*AP,AP*) *MHC* genotype had the lowest risk (*OR=*0.13) relative to the (*0,0*) *MHC* genotype of any that we identified (Table 4; Fig.1). Therefore, this group was used to normalize the MHC haplotype risk effects. We show that all *MHC* genotypes, except (*c1*), are intermediate between the two causal models (Fig. 4). By contrast, for (*c1,c1*) and (*c1,ER*) genotypes, the observations exceed the expectations of both models (Fig. 4).

When the other non-*MHC* “risk” loci are included in the analysis, observations are closer to the additive model. Thus, the estimates from a multiplicative model exceed observations by1-2 orders of magnitude (Figs. 5-7). As demonstrated previously for a different definition of the (*G*) subset [3], the distribution of penetrance values in the general population (*Z*) is incompatible with a lognormal distribution [3] - i.e., the distribution expected for a multiplicative model. In the present iteration of the model, defining the set (*G*) to include all genotypes, which that have a non-zero expected penetrance, the bimodality of the distribution can be established with certainty (*File S1*). Consequently, based upon both theory and observation, a multiplicative model for the accumulation of genetic risk in MS is inappropriate.

The additive model, in general, performed better in these circumstances (Figs. 4-7). Nevertheless, it does not explain perfectly the accumulation of genetic risk in MS. First, (*c1*) CEH genotypes consistently exceed the additive expectations (Fig. 4). Second, effect of a given *MHC* haplotype is dependent on its companion *MHC* haplotype in a genotype (Fig. 2). Third, the effect of the 3 non-*MH*C “risk” haplotypes is not consistent across all *MHC* genotypes (Fig. 3). And fourth, when more loci are included in the analysis, the observations become increasingly less than what is predicted by the additive model (Figs. 5-7). Taken together, these lines of evidence indicate that the accumulation of genetic risk from these “susceptibility loci” is inconsistent with both an additive and a multiplicative model. Rather, the magnitude of any change in disease-risk associated with the inclusion of additional “susceptibility loci” seems to depend upon the exact state at each “risk-locus” and on the interaction across all loci. Such a conclusion is also anticipated on the basis of theoretical considerations (*File S1*).

The *MHC* is known to have a remarkable diversity [63]. In the WTCCC population, there were 29 *HLA-A* alleles, 29 *HLA-C* alleles, 55 *HLA-B* alleles, 35 *HLA-DRB1* alleles, and 16 *HLA-DQB1* alleles. Moreover, these alleles did not exist in isolation but, rather, as part of 10,078 unique CEHs, 810 of which accounted for 71% of all the CEHs present in the WTCCC dataset [26]. Also, even if some CEHs share common features, such as carrying the (*H+*)*-*haplotype, the degree of association with MS varies depending upon the exact CEH considered (*S2 File; Tables S2 & S3*). For example, both (*c2*) and (*c3*) CEHs carry the (*H+*)-haplotype, but their MS-association differed significantly (*z=4.8; p<10*^*-6*^). It might be tempting to attribute this difference to (*c3*) carrying the potentially “protective” *HLA-A*02:*01 allele (*S2 File; Table S2*). However, other *HLA-A*02:*01 and (*H+*) carrying CEHs (e.g., *c50, c58*, and *c139*) do not seem to be similarly protected (*S2 File; Table S2*). Finally, each identified CEH probably represents a diverse set of CEHs. Thus, because the 3 mb genomic region from *HLA-A* to *HLA-DQB1* is quite “gene-dense”, each of the CEHs that we defined, almost certainly, represent groups of CEHs, which carry many other linked polymorphisms.

Although the non-*MHC* “risk” regions used for this analysis are likely to be less variable than the MHC, these regions span large amounts of DNA (200-680 kb) and they generally have hundreds of highly conserved SNP-haplotypes across each region. Moreover, despite the fact that authors sometimes identify specific genes as being MS-associated [13,14], the truth is that we have no basis for deciding which gene or genes within a region are responsible for the association. We cannot exclude the possibility that, within these regions, as within the MHC, there might exist “risk” or “protective” alleles interacting with each other. If so, the likelihood that any simple probability model (either additive or multiplicative) will adequately describe genetic-susceptibility to MS seems quite remote.

However, such complexity fits well with the model of genetic-susceptibility presented in the *Introduction* and more fully developed in the *S1 File*. Thus, MS-susceptibility – i.e., membership in the subset (*G*) – seems to be confined to a small subset (~2.2– 4.5%) of the general population (*File S1; Table S1*) and, yet, this susceptibility is a prerequisite to getting MS, with members of the (*G-*) subset having no chance of getting MS, regardless of what environmental experiences they have (*S1 File*). Moreover, despite the fact that the Class II (*H+*)-haplotype is, by far, the strongest, and most significant, MS-associated genetic factor (p<<10^-300^) of any in the genome and has been known for over a half a century [11,15-22,26], only a tiny fraction of (*H+*) carriers are even “susceptible” to getting MS (*File S1*). This observation indicates that at least with respect to the (*H+*)-haplotype, “genetic-susceptibility” to MS requires the combined effects of different genes (*File S1*). The presence of (*H+*), by itself, does not increase disease-risk.

Nevertheless, despite the necessity of being a member of the (*G*) subset in order for a person to develop MS, environmental factors are also required. Indeed, once “genetic-susceptibility” is established in an individual, these environmental factors, together with certain stochastic processes, are entirely responsible for determining who does and who does not ultimately develop MS (*File S1*). Some of these causal environmental factors seem to occur *in utero* or, possibly, in the early post-natal period; others seem to occur during adolescence; and still others seem to occur later [50,51,65-67]. There is strong evidence that Epstein Barr viral infections (especially those associated with symptomatic infectious mononucleosis) are causally associated. There is also strong circumstantial evidence that Vitamin D deficiency is an important factor [50,51,65-67]. Other factors (e.g., smoking, obesity, and possibly other infections) may also play a role [50,51,65-67]. Regardless of the identity and role of each factor, however, it seems that, collectively, these environmental events (which currently occur as “population-wide” exposures) are major determinants of whether or not the disease will develop in a “susceptible” individual (*File S1*). For example, although the (*H+*)-haplotype, as noted, has the strongest association with MS of any, the possession of this haplotype seems only to make (*G*) subset membership more likely but does not seem to alter the likelihood of actually getting MS once (*G*) subset membership is established (Table 1; *S1 File*).

Similarly, the well-known gender-bias in MS prevalence seems to be largely explained by an increased responsiveness of “susceptible” women to the environmental factors involved in MS pathogenesis (*File S1*). Nevertheless, men are more likely to be members of the (*G*) subset than are women (*File S1*). Such a finding might seem surprising given the facts that 1) all of the ~200 MS-linked loci are on autosomal chromosomes [13,14]; 2) association studies specifically focused on the *X*-chromosome have not suggested the presence of any *X*-linked associated loci [7]; and, finally, 3) it is hard to rationalize how women could possibly be more or less likely to possess any specific autosomal genotype compared to men (*S1 File*). Indeed, in the WTCCC, women seemed to be equally likely to possess both the “risk” and the “protective” CEH combinations compared to men. Nevertheless, if (*G*) subset membership depends upon specific genetic combinations, and defining the subset (*G*_*a*_) to represent the autosomal genotypes in the general population (*Z*), it is certainly plausible that men and women could be equally likely to possess each of its members and, yet, for any specific (*k*^*th*^) member with genotype (*G*_*ak*_), it could well be the case that:

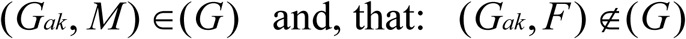

Such a circumstance would represent another example of “genetic-susceptibility” to MS requiring the combined effect of different genetic traits (*File S1*).

Moreover, from the epidemiological observations regarding changes in sex-ratio that have taken place over time [68], the response curves for “susceptible” men and women to increasing levels of environmental exposure can be derived quantitatively (*Supplemental Fig. C; File S1*) and it is clear from these response curves that women are, indeed, more responsive (probably physiologically) than men to the causal environmental factors involved in MS pathogenesis [3,4,68]. Moreover, this analysis strongly suggests that the relevant environmental exposures must be occurring currently at “population-wide” levels (*S1 File*). Such a conclusion is fully consistent with the same conclusion reached from observational studies in adopted individuals, in siblings and half-siblings raised together or apart, in conjugal couples, and in brothers and sisters of different birth order, which have generally indicated that MS-risk is unaffected by the childhood or other micro-environments [69–75].

By contrast, comparing the penetrance in the (*H+*) and (*H-*) subsets of *MZ*-twins, the fact that there is little difference in penetrance between the (*H* +,*G, IG*_*MS*_) and (*H* –,*G, IG*_*MS*_) subsets strongly suggests that there is also little difference in penetrance between the (*H* +,*G*) and (*H* –,*G*) subsets. Indeed, as demonstrated in *File S1*, despite the fact that the disease association for the (*H+*) subset is, by far, the strongest and most significant of any in the entire genome [11,15-22], this association is due mostly to the fact that: *P*(*G* | *H* +) ≫ *PG* | *H* –).

In the study of human genetics there has been a long-running debate between the so-called “common-disease, common variant” (*CDCV*) and the “common-disease, rare variant” (*CDRV*) hypotheses [76]. Nevertheless, with our improved genetic sophistication, it has become increasingly clear that, in different specific circumstances, either (or both, or neither) hypotheses could be operative [76]. In fact, our observations also support this notion. For example, on the one hand, all of the *MHC* CEH combinations, which impact MS-susceptibility, are quite rare. None has a population frequency in controls of more than 6.2% and the large majority of them have population frequencies well below 1% (*S2 File; Tables S2&S3*). On the other hand, considered collectively, those CEH combinations, which include the Class II (*H+*)-haplotype, have a WTCCC Control population frequency of 23%. Indeed, this particular haplotype-group is the most prevalent (and, therefore, the most highly selected) of all such Class II haplotype combinations in northern population [26].

Consequently, regardless of whether one considers these observations to be supportive of an association between MS-susceptibility and “common” or “rare” variants, the fact remains that, whether considered individually or collectively, the most prevalent, and therefore the most highly selected [26], CEHs are those that are also associated with the highest MS-risk (*S2 File; Tables S2&S3*). Thus, it is clear that these particular CEHs must come with both adaptive and deleterious consequences for the individual. Also, although the CEH composition differs markedly among long-separated human populations (*File S2; Tables 4a & 4b*), specific CEHs are still being strongly selected in each of them [26]. Consequently, the benefits of the adaptive features of these CEHs must outweigh the risk of any deleterious ones. Obviously, for circumstances, either in which the risk of MS is small or in which MS has little impact on an individual’s eventual number of surviving children, even a modest advantage in favor of a specific CEH might still cause it to be selected. In this regard, a recent French study estimated that women with MS had 31% fewer children than their contemporary controls [77]. If this observation is correct, it suggests that there is a strong selective disadvantage to having MS. Therefore, the explanation for the benefits of these MS-associated CEHs outweighing the risks is likely to lie in an individual’s low risk of MS rather than the disease having little impact on their fertility. Based on our observations, this seems likely to be the case. Thus, because natural selection can only select against those genotypes, which actually carry risk (relative to other genotypes), the fact that so few members of the “susceptible” (*G*) subset ever actually develop MS makes such a favorable tradeoff between adaptive and deleterious features considerably more likely to occur.

Our results also bear on the common notion that there is a considerable amount of “missing heritability” in both MS and other complex genetic disorders [78-80]. First, as discussed in *File S1*, much of the variability in MS expression (even among “genetically-susceptible” individuals) is attributable to stochastic processes that are unrelated to either environmental or genetic factors. Second, MS expression is related to an interaction between the environmental and genetic factors involved in MS pathogenesis; neither alone are sufficient and both are necessary (*File S1*). Third, both theoretically (*File S1*) and observationally (Figs 2 & 3) specific gene-gene combinations are crucial determinants of “susceptibility” to MS - a circumstance which renders the common (additive) methods of estimating heritability unreliable [77]. And fourth, with over 200 independent MS-associated genetic regions [5-14], each potentially with more than one “susceptible state” (e.g., the MHC), there are so many possible combinations of states at these loci that, almost certainly, every person with MS possesses a unique combination. If, as indicated by our results, only a few of these combinations are members of the (*G*) subset, even of combinations that are similar to each other (*File S1* & *File S2; Table S2*), then there are more than enough genetic associations identified already to account fully for membership in the (*G*) subset. Naturally, many more MS-associated loci may yet be identified in the future although their existence is not necessary (*File S1*).

Alternatively, if “missing heritability” is meant to imply only that our genetic model cannot predict accurately the occurrence of MS, then it is true that a substantial amount of the “heritability” remains unexplained. Indeed, the environmental factors, gene-gene combinations, gene-environment interactions, and stochastic factors, which underlie the development of MS in any individual, are poorly understood, thereby making any accurate prediction of MS occurrence, at present, impossible.

Finally, it is worth noting that the nature of “genetic susceptibility” developed in this manuscript is applicable to a wide range of other complex genetic disorders such as type-1 diabetes mellitus, Crohn’s disease, ulcerative colitis, and rheumatoid arthritis. Indeed, base solely upon *Proposition #1* (*S1 File*), any disease for which the MZ-twin concordance rate is substantially greater than the life-time risk in the general population, only a small fraction of the population can possibly be in the “genetically susceptible” subset (i.e., have any chance of developing the disease). Moreover, any disease for which the MZ-twin concordance rate is substantially less than 100% must, in addition to “genetic susceptibility”, include environmental factors, stochastic factors, or both in the causal pathway leading to the disease.

## Materials & Methods

### Ethics Statement

This research has been approved by the University of California, San Francisco’s Institutional Review Board (IRB) has been conducted according to the principles expressed in the Declaration of Helsinki.

### Study Participants

#### Wellcome Trust Case Control Consortium (WTCCC)

This multinational study cohort consists of 18,872 controls and 11,376 cases with MS and has been described in detail previously [13,14]. However, SNP haplotype data was unavailable for 380 controls and 232 cases. Of the cases, 72.9% were women, the average age-of-onset was 33.1 years, and the mean Extended Disability Status Score (EDSS) was 3.7 [14]. The patients enrolled in this study (except for a few African Americans from the United States) were of European ancestry. The large majority (89%) of the cases had a relapsing-remitting onset [13]. The diagnosis of MS was made based upon internationally recognized criteria [81-83]. Control subjects were composed of healthy individuals with European ancestry [13]. The Ethical Committees or Institutional Review Boards at each of the participating centers approved the protocol and informed consent was obtained from each study participant. The WTCCC granted data access for this study.

### Genotyping, and Quality Control

The genotyping methods and quality control for the WTCCC have been described in detail previously [13,14,16,18,19]. All genotyping was performed on the Illumina Infinium platform at the Wellcome Trust Sanger Institute. Case samples were genotyped using a customized Human660-Quad chip. Common controls were genotyped on a second customized Human1M-Duo chip (utilizing the same probes). After quality control, this provided data on 441,547 autosomal SNPs scattered throughout the genome in both MS patients and controls. The identities of the five HLA alleles in the MHC region (*A, C, B, DRB1* and *DQB1*) were determined for each participant by imputation using the HIBAG method [84].

### Statistical Methods

#### Phasing

Both the phasing of alleles at each of five *HLA* loci (*HLA-A, HLA-C, HLA-B, HLA-DRB1* and *HLA-DQB1*) and the phasing of the SNP-haplotypes surrounding the Class II region of the *DRB1* gene were accomplished using previously published probabilistic phasing algorithms [23,24,85,86]. SNP-haplotypes from 3 of the 102 non-MHC genomic regions, which had been identified previously as being significantly MS-associated, were also included in our analysis [24]. In our previous report, the MS-associated SNP haplotypes were numbered (arbitrarily) from 1 to 932. These three particular regions (arbitrarily labeled *d1, d2*, and *d3*) were selected based on their having a “risk” SNP-haplotype with 500 or more representations in the WTCCC dataset and also having the largest *ORs* for disease-association of any haplotype meeting this specification. The reason for choosing only three regions was that, when more regions were added, there were an insufficient number observations to estimate the *ORs* for any of the possible higher order combinations. These three regions were located at chromosomal locations 3p24.2, 14q24.1, and 16p13.13 and in the vicinity, respectively, of the genes EOMES, ZFP36L1, CLEC16A [13,14]. Chromosome 3; Region 22 (*d1*) spanned 0.65 mb of DNA and the 11-SNP-haplotype (number 234) was used [24]. Chromosome 14; Region 78 (*d2*) spanned 0.68 mb of DNA and the 3-SNP-haplotype (number 734) was used [24]. Chromosome 16; Region 85 (*d3*) spanned 0.20 mb of DNA and the SNP-haplotypes (numbers 814, 818, and 822) were combined into a single 15-SNP haplotype [24]. This was done because each of these risk-haplotypes were adjacent to each other and because the individual risk SNP haplotypes were part of the same extended 15-SNP-haplotype.

#### Haplotype Frequencies and Association Testing

Disease association tests, as measured by *ORs* and confidence intervals (*CIs*), were calculated for each of the CEHs and each of the 3 non-MHC risk haplotypes either alone or in different combinations. The WTCCC data was considered in its entirety and not further stratified. MS-associated haplotypes were analyzed by grouping them into five categories of CEHs, which consisted of: 1) (*H+*)-carrying CEHs (i.e. those containing the *HLA*-*DRB1*15:01~HLA*-*DQB1*06:02~a1* haplotype, *S2 File and Table S2*; other increased risk or “extended risk” (*ER*) CEHs (*c23, c27, c34, c46, c68, c81, c85, c96*, and *c107*) as shown in *S2 File* (*Table S3*); 3) decreased risk or “all protective” (*AP*) CEHs (*c5, c15, c18, c24, c30, c32, c51*, and *c73*) as shown in *S2 File* (*Table S3*); 4) all CEHs not in the (*H+*), (*ER*), or (*AP*) groups (*0*) CEHs; and 5) the (*c1*) CEH by itself. We also explored a “protective” group, which excluded the (*c5*) CEH. However, this analysis is not presented because the findings were the same as when the *AP* group was analyzed as a whole. In many circumstances, an individual’s *MHC* genotype was specified by the haplotype combination that they possessed. For example, by this convention, an individual homozygous for (*H+*) would be characterized as having the (*H+,H+*) *MHC* genotype. By contrast, a heterozygous individual would be characterized as having the (*H+,0*), the (*H+,ER*), the (*H+,c1*), or the (*H+,AP*) *MHC* genotype In the principal analysis, all MS-associations were assessed compared to a reference group consisting of the (*0,0*) *MHC* genotype. Similarly, when the disease associations for those “non-risk” CEHs carrying the *HLA-DRB1*03:01~HLA-DQB1*02:01~a6* haplotype were assessed, other carriers of this haplotype were also excluded from the (*0,0*) reference group. For notational simplicity, when using the (*AP,AP*) MHC genotype as a reference, this genotype was referred to as (*AP\**).

Disease associations for the risk SNP-haplotypes on Chromosomes 3, 14, and 16, were assessed compared to a reference group consisting of the (*0,0*) *MHC* genotype, and excluded individuals carrying their risk-haplotypes at these chromosomal locations. We designate (collectively) all non-risk-haplotypes at each of these chromosomal locations as the (*0*) haplotype at each locus.

Significance of the differences between two *ORs* in disease association for any haplotype or haplotype combination was determined by z-scores calculated for the differences in the natural logarithm of the *ORs* for any two haplotypes. As discussed earlier, pair-wise comparisons of *ORs* are independent of the reference group chosen. The *MHC* genotype (*0,0*) had the largest sample size of any and, therefore, in order to maximize the statistical power to detect differences, the *ORs* used for pair-wise comparisons within the *MHC* were estimated relative to a reference group consisting of the (*0,0*) genotype at both the *MHC* and also at the any non-MHC locus included in the comparison. As noted in the *Introduction*, such a method eliminates the common reference group disease-risk to yield an estimate of the pairwise *RR*. Within the WTCCC cohort, we used a principal components (*PC*) analysis excluding *MHC* SNPs (Eigensoft) to correct the observations in *Tables S2 & S3 (S2 File)* for the possible effects of population stratification, as well as regression analysis to correct for the possible effects of geographic heterogeneity [25]. These adjustments did not significantly alter any of the associations shown in (*S2 File; Tables S 2& S3*).

#### Evaluating Additive and Multiplicative Risk-models

The *ORs* for the *MHC* alleles (*H+, ER*, and *0*) were determined relative to the (*AP*^*\**^) reference group, which was assigned a value of (*R*_*b*_*=R*_*AP\**_*=*1). These observed *ORs* were used to estimate the *RRs* associated with each set of *MHC* alleles and, in turn, these *RRs* were used to assess the appropriateness of the additive and multiplicative risk-models for the different allelic combinations at the MHC. Subsequently, using a reference group consisting of the (*0,0*) *MHC* genotype, we determined the *ORs* for susceptibility alleles in the three non-MHC susceptibility regions – (*d1*), (*d2*) and (*d3*). The (*0,0*) *MHC* genotype was chosen as the reference because there were too few representations of the (*AP,AP*) *MHC* genotype in the WTCCC dataset. Nevertheless, these observed *ORs* were mathematically converted into *ORs* relative to the (*AP,AP*) *MHC* genotype and these re-referenced *ORs*, together with the *ORs* actually observed for the different allelic combinations at the *MHC*, were used to estimate the *RRs* associated with each allelic combinations at these four genomic locations (the *MHC* plus the three non-*MHC* susceptibility regions). These estimated *RRs* were then used to assess the appropriateness of the additive and multiplicative risk-models for the different allelic combinations of these four susceptibility regions. In all cases, only *ORs* estimated from combinations with *≥*15 representations in the WTCCC were considered.

## Acknowledgements

None.

## Parameter Definition

## Supporting Information Captions

***File S1***. This *Supplemental File* describes, in detail, the susceptibility Model used for this manuscript and rigorously develops its logical framework. It includes the methods used to estimate both the “population” parameters such as:

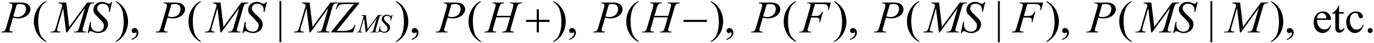

as well as the “non-population” parameters of interest such as:

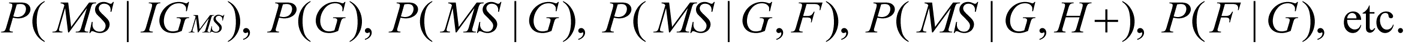

(See *Table 1, Main Text* for the definitions of the different parameters used in the Model)

It also describes, in detail, the manner by which the values of these non-population parameters can be estimated from the directly observable population parameters.

***File S2***. This *Supplemental File* describes composition of the *CEHs* found in the WTCCC dataset as well as their individual relationships to MS susceptibility and also how this *CEH* composition differs between populations around the world.

Furthermore, this file considers the theoretical underpinnings for the commonly used additive and multiplicative Models for the accumulation of disease “risk” with increasing number of “risk haplotypes” being present in an individual’s genotype

